# Chloride-dependent mechanisms of multimodal sensory discrimination and neuropathic sensitization in *Drosophila*

**DOI:** 10.1101/2021.12.10.472093

**Authors:** Nathaniel J. Himmel, Akira Sakurai, Jamin M. Letcher, Atit A. Patel, Shatabdi Bhattacharjee, Maggie N. Benson, Thomas R. Gray, Gennady S. Cymbalyuk, Daniel N. Cox

## Abstract

Individual sensory neurons can be tuned to many stimuli, each driving unique, stimulus-relevant behaviors, and the ability of multimodal nociceptor neurons to discriminate between potentially harmful and innocuous stimuli is broadly important for organismal survival. Moreover, disruptions in the capacity to differentiate between noxious and innocuous stimuli can result in neuropathic pain. *Drosophila* larval Class III (CIII) neurons are peripheral noxious cold nociceptors and gentle touch mechanosensors; high levels of activation drive cold-evoked contraction (CT) behavior, while low levels of activation result in a suite of touch-associated behaviors. However, it is unknown what molecular factors underlie CIII multimodality. Here, we show that the TMEM16/anoctamins *subdued* and *white walker* (*wwk*; *CG15270*) are required for cold-evoked CT, but not for touch-associated behavior, indicating a conserved role for anoctamins in nociception. We also evidence that CIII neurons make use of atypical depolarizing chloride currents to encode cold, and that overexpression of *ncc69*—a fly homologue of *NKCC1*—results in phenotypes consistent with neuropathic sensitization, including behavioral hypersensitization and spontaneous nociceptor activity, making *Drosophila* CIII neurons a candidate system for future studies of the basic mechanisms underlying neuropathic pain.

## INTRODUCTION

Noxious stimuli are transduced by high-threshold sensory neurons referred to as nociceptors, and these sensory neurons often respond to more than one stimulus type – a property called sensory poly- or multimodality ^1-5^. Although the ability to differentiate between sensory modalities is inarguably important for organismal survival, it remains relatively poorly understood what molecular mechanisms facilitate sensory multimodality within single neurons or neural subtypes; and these systems are of direct importance human health, as the inability to discriminate between noxious and innocuous stimuli is thought to underlie chronic neuropathic pain ^6-8^.

Like other animals, *Drosophila melanogaster* can sense and respond to noxious stimuli. In larvae, nociception primarily begins in peripheral dendritic arborization neurons of the Class III (CIII) and Class IV (CIV) subtypes. CIV neurons are polymodal high temperature, mechanical, and chemical nociceptors, and activation of class IV neurons by any one of these sensory modalities elicits a corkscrew-like rolling behavior ^9-13^. In contrast, CIII neurons are multimodal cold nociceptors and gentle-touch mechanosensors, and activation of CIII neurons drives stimulus-specific behaviors: acute noxious cold primarily elicits a highly stereotyped, bilateral contraction (CT) response, wherein the body rapidly shortens along the head-to-tail axis ^14^, while gentle touches elicit a suite of behaviors, including head-withdrawal, head casting/turning behavior, and reverse locomotion ^15^.

CIII neurons function via a high-low threshold detection system, whereby high levels of CIII activation (and strong Ca^2+^ transients) are associated with CT, and low levels of activation (and relatively modest Ca^2+^ transients) with touch-behaviors. This is evidenced in Ca^2+^ imaging experiments and via optogenetics, where strong optogenetic activation of CIII neurons elicits CT and less strong activation primarily drives head withdrawals ^14^. However, molecular factors underlying this high-low threshold filter have not been elucidated. Given the relatively constrained function and modality-specific behavioral outputs of CIII, this system constitutes a good target for elucidating mechanisms underlying multimodality in single neural classes.

Stimulus-evoked CIII calcium transients are thought to occur as a result of the activation of Transient Receptor Potential (TRP) channels, a superfamily variably selection cation channels which participate in nociception across a wide variety of species ^16^. In *Drosophila*, the TRP genes *Pkd2, NompC*, and *Trpm* are required for cold nociception and gentle touch mechanosensation^14^. In vertebrates, Ca^2+^-dependent channels of a variety of subtypes (*e*.*g*. anoctamin/TMEM16 channels ^17^) interact with TRP channels in nociceptive systems in order to drive appropriate levels of neural activation, making them a potential candidate mechanism underlying cold and touch discrimination. It has been previously shown that the *Drosophila* gene *subdued* encodes a Ca^2+^-activated Cl^-^ channel (CaCC) ^18,19^, and that it is necessary for high temperature-evoked rolling ^20^. However, it is unknown whether or not subdued might function to encode stimulus-specific sensory information in multimodal neurons; we therefore questioned whether anoctamins might function in CIII in a modality-specific fashion.

In the present study we tested the hypothesis that anoctamins expressed in CIII are selectively required for cold nociception. We have found that the anoctamins *subdued* and *white walker* (*wwk*; *CG15270*)—here shown to be orthologous to human ANO1/2 and ANO8, respectively—are required for cold nociception. Interestingly, anoctamins participate in an excitatory capacity, suggesting that CIII neurons make use of excitatory Cl^-^ currents, and we provide additional evidence that CIII neurons likely use depolarizing Cl^-^ currents to selectively encode cold. Further, we demonstrate that overexpression of *ncc69* (a fly homologue of *NKCC1*) is sufficient for driving a neuropathic pain-like state in *Drosophila* larvae – a phenotype mechanistically analogous to human neuropathic pain following spinal cord injury.

## MATERIALS AND METHODS

### Fly Strains

All *Drosophila melanogaster* stocks were maintained at 24°C under a 12:12 light cycle. Genetic crosses were raised at 29°C under a 12:12 light cycle in order to accelerate development. 3^rd^ instar larvae were used for all experiments. Publicly available transgenic strains were originally sourced from the Bloomington Drosophila Stock Center (B) and the Vienna Drosophila Resource Center (v) and included: *T2A-GAL4*^*wwk*^ (B76649); *UAS-Aurora* (B76327); *UAS-mCD8::GFP* (B5130); *wwk*^*MI03516*^ (B36976); *Df(2L)b87e25* (*wwk df*, B3138); *subdued*^*MI15535*^ (B61082); *Df(3R)Exel6184* (*subdued df*, B7663); *UAS-subdued RNAi* (v32472 and v108953); *UAS-wwk RNAi* (B28650 and B62282); *UAS-ncc69 RNAi* (B28682); and *UAS-kcc RNAi* (B34584). *UAS-kcc* was provided by Dr. Mark Tanouye; *UAS-ncc69* was provided by Dr. Don van Meyel; *UAS-subdued* and *GAL4*^*c240*^ were provided by Dr. Changsoo Kim. *GAL4*^*19-12*^ is available upon request.

*CIII::tdTomato* and *UAS-wwk* were developed for this study and are available upon request. For *GAL4*^*CIII*^*::tdTomato* we utilized Gateway cloning to combine the *R83B04* (CIII) enhancer with *CD4-tdTomato* by LR reaction (Invitrogen ref: 12538-120). Transgenic flies carrying a 2^nd^ chromosomal insertion of *CIII:tdTomato* (docking site: VK37) were generated by Genetivision. The *R83B04* enhancer-containing entry vector was provided by the FlyLight Project. *pDEST-HemmarR2*—the vector for enhancer driven *CD4-tdTomato* expression—was provided by Dr. Chun Han. *For UAS-wwk*, full-length cDNA was synthesized (GenScript) and subcloned into *pUAST-attB*. Transgenic *UAS-wwk* flies carrying a 3^rd^ chromosomal insertion of *UAS-wwk* (docking site: VK20) were generated by Genetivision.

### Class III isolation and qRT-PCR

Isolation of CIII neurons followed a previously described protocol ^21^. In brief, 40-50 third instar larvae with mCD8::GFP-tagged CIII neurons were collected and washed in ddH_2_0, dissected using microdissection scissors, and then dissociated in PBS using a glass dounce. CIII neurons were then isolated using superparamagnetic beads (Dynabeads MyOne Steptavidin T1, Invitrogen) that were conjugated to biotinylated anti-mCD8a antibody (eBioscience). RNA was isolated from these neurons using the RNeasy Mini Kit (Qiagen) and qRT-PCR analysis was performed in triplicate using pre-validated Qiagen QuantiTect Primer Assays for *CG15270* (QT00936754) and *subdued* (QT00978131). Data are reported as C_q_ value ± standard deviation.

### Cold Nociception Assay

3^rd^ instar larvae were raised in and collected from standard fly food vials, gently washed in water, then acclimated to a room temperature, moistened, black-painted aluminum arena. Once locomotion resumed, the arena was transferred to a prechilled Peltier device (TE Technologies, CP0031) under the control of a thermoelectric temperature controller (TE Technologies, TC-48-20). Behavior was recorded from above by a Nikon D5200 DSLR camera, and the first 5 seconds of footage following contact between the arena and Peltier plate were used for behavioral analysis. Videos were converted to the avi format by Video to Video software (https://www.videotovideo.org/), and then further processed in the FIJI distribution of ImageJ. Virtual stack images of larvae were converted to grayscale, thresholded, and subsequently skeletonized into single pixel-wide lines representing larval length from tip to tip. Length measurements were normalized to the length at time 0, and CT was counted if the larvae passed the CT threshold at any point during the analysis. The CT threshold was determined as in previous studies, as the average peak *w*^*1118*^ decrease in length + 1.5x the standard deviation; this resulted in a CT threshold (here called strong CT) of approximately 0.7 (also presented as -0.3 change in normalized body length or 30% reduction in body length). Data are reported as % CT and the peak magnitude of decrease in larval length.

### Mechanosensation Assay

3^rd^ instar larvae were collected, gently washed, then acclimated to the same arena used in cold plate assays. Mechanosensitivity was scored similarly to previous reports ^14,15^: in brief, animals were brushed on an anterior segment by a single paintbrush bristle and behavior was observed through a Zeiss Stemi 305 microscope. Gentle touch behaviors include pausing/hesitating, head/anterior withdrawal (AKA hunching), head casting/turning, and reverse locomotion. Each subject was given 1 point if it performed one of these behaviors (for a maximum of 4 points per trial). Each animal was subject to three touch trials with 30s intervals between each trial. The scores from the three trials were summed (for a maximum of 12 points per subject), and averaged across each genotype.

### Optogenetics

3^rd^ instar larvae were collected, gently washed, then acclimated to a glass plate. The plate was then transferred to a custom-built optogenetics-behavior rig fitted with bottom-mounted blue LED illumination and a top-mounted Canon Rebel T3i DSLR camera. Blue light, video recording, and larval tracking was handled automatically by Noldus EthoVision software; behavior was recorded for 5 seconds in the absence of blue light, for 10 seconds with blue light illumination, and for 5 additional seconds in the absence of blue light. Larval behavior was quantified by automated tracking of larval area from above.

### Microscopy

For morphometric analyses, GFP tagged CIII neurons in 3^rd^ instar larvae were imaged on a Zeiss LSM 780 confocal microscope at 200X magnification. Images were collected as z-stacks with a step size of 2.0 µm and at 1024×1024 resolution. Maximum intensity projections of z-stacks were exported using Zen (blue edition) software and analyzed using the Analyze Skeleton ImageJ plugin. Metrics were compiled using custom Python algorithms freely available upon request.

To image *GAL4* expression patterns, larvae were likewise imaged under Zeiss LSM 780 confocal microscope at various magnifications (scale bars present on relevant images), at various locations (indicated in relevant figures/legends). Images were viewed and exported from Zen (black edition) software and labeled in Adobe Illustrator.

### Electrophysiology

Demuscled fillet preparations were made from 3^rd^ instar larvae were placed on the bottom of an experimental chamber (200 µL) filled with HL-3 saline, which consisted of (in mM) 70 NaCl, 5 KCl, 1.5 CaCl_2_, 20 MgCl_2_, 10 NaHCO_3_, 5 trehalose, 115 sucrose, and 5 HEPES (pH 7.2). For the low-chloride saline, sodium chloride and magnesium chloride were replaced with sodium sulfate (Na_2_SO_4_, 35 mM) and magnesium sulfate (MgSO_4_, 20 mM). The osmolarity of the low-chloride saline was adjusted to become equal to HL-3 saline (355-360 mOsm) by adding sucrose (up to 170 mM in total). The specimen was constantly perfused with HL-3 saline (30-40 µL/sec). Switching between HL-3 and low-chlorine saline was done by a three-way stopcock attached to the inlet tube.

Cold temperature stimulation was administered by passing the saline through an in-line solution cooler (SC-20, Warner Instruments, Hamden, CT, USA) connected to the controller device (CL-100, Warner Instruments, Hamden, CT, USA). The saline temperature was constantly monitored by a thermometer probe (BAT-12, Physitemp, Clifton, NJ, USA) placed adjacent to the preparation.

The spiking activity of a GFP-tagged CIII neurons was recorded extracellularly by using a borosilicate glass micropipette (tip diameter, 5–10 µm) by applying gentle suction. The electrode was connected to the headstage of a patch-clamp amplifier (MultiClamp 700A, Molecular Devices, San Jose, CA, USA). All recordings were made from ddaA in the dorsal cluster of the peripheral sensory neurons. The amplifier’s output was digitized at 10 kHz sampling rate using a Micro1401 A/D converter (Cambridge Electronic Design, Cambridge, UK) and acquired into a laptop computer running Windows 10 (Microsoft, Redmont, WA, USA) with Spike2 software v. 8 (Cambridge Electronic Design, Cambridge, UK). Bursting spikes were identified as those with inter-spike intervals of less than 0.2 seconds.

### Phylogenetics

Starting with previously characterized ANO sequences from human (NCBI CCDS), mouse (NCBI CCDS), and *D. melanogaster* (FlyBase), an ANO sequence database was assembled by Blastp against protein sequence/model databases at Ensembl and the Okinawa Institute of Science and Technology Marine Genomes Unit. In order to maximize useful phylogenetic information, only Blast hits >300 amino acids in length with an E-value less than 1E−20 were retained.

Next, CD-HIT was used to identify and remove duplicate sequences and predicted isoforms (threshold 90% similarity), retaining the longest isoform to maximize phylogenetic information. Phobius was then used to predict TM topology; in order to further maximize phylogenetic information, sequences with fewer than five predicted TM segments were removed.

Sequences were thereafter aligned via MAFFT using default settings. The final phylogenetic tree was generated via IQ-Tree by the maximum likelihood approach, using an LG+R7 substitution model (as determined by ModelFinder). Branch support was calculated by ultrafast bootstrapping (UFboot, 2000 replicates). In order to identify duplication events, the maximum likelihood phylogeny was reconciled using NOTUNG 2.9.1. In order to formulate the most parsimonious interpretation of the resulting trees, weakly supported branches were rearranged (UFboot 95 cutoff) in a species-aware fashion against the species cladogram in **Figure 4A**. Edge weight threshold was set to 1.0, and the costs of duplications and losses were set to 1.5 and 1.0, respectively. Trees were visualized using iTOL and Adobe Illustrator.

### Statistical Analyses

Due to a growing call for statistical analyses that do not rely on *p*-values ^22-30^, differences in mean and population proportions were analyzed using both traditional frequentist statistics and Bayesian alternatives.

Population proportions are presented as % ± standard error of the proportion (SEP); differences in proportion were assessed by one-tailed (for 5°C experiments) or two-tailed (for 10°C experiments) z-test with a Bonferroni correction and the Bayesian A/B Test. All other measures are presented as mean ± standard error of the mean (SEM) unless otherwise noted; differences were assessed by frequentist one- or two-way ANOVA with Dunnett’s *post-hoc* tests and Bayesian equivalents^31^, or frequentist and Bayesian t-test (in the case of comparisons between only 2 groups). Electrophysiology low Cl^-^ wash experiments were performed within-subjects. For some subjects recordings were lost during the low Cl^-^ or washout phase; to account for these missing data, differences were assessed using a linear mixed model and Bayesian repeated measures ANOVA (both analyzing the effect of different washes on firing frequency). z-tests were performed in Microsoft Excel, two-way ANOVA in GraphPad PRISM, and all other analyses in JASP (Bayesian analyses using default prior probability distributions or priors adjusted by the Westfall-Johnson-Utts method) ^32^.

For frequentist analyses, statistical significance as assessed at α=0.05. For Bayesian analyses, degree of support for rejecting the null hypothesis was inferred by computed Bayes Factors (BF), via a modification on the method originally proposed by Jeffreys ^33^: “null hypothesis supported” (BF_10_ < 1); “weak” (BF_10_: 1-3); “substantial” (BF_10_: 3-10); “strong” (BF_10_: 10-30); “very strong” (BF_10_: 30-100); “decisive” (BF_10_ > 100).

All graphs were generated using GraphPad PRISM (GraphPad Software, La Jolla, California, USA) and Adobe Illustrator. Relevant n, adjusted p, and BF_10_ values are listed in figure legends. In figures, asterisks (*) indicate statistical significance at α=0.05; daggers (†) indicate weak evidence in favor of the alternative hypothesis; double daggers (‡) indicate at least substantial evidence in favor of the alternative hypothesis; and families of comparisons are grouped by overhead bars.

## RESULTS

### The anoctamins *subdued* and *CG15270* are required for cold nociception

Previous work has demonstrated that TRP channels are required for both CIII-dependent cold nociception and gentle touch mechanosensation^14^; however, no genes have been identified which are selectively required for CIII cold nociception. As vertebrate TRP channels function alongside anoctamin/TMEM16 channels in vertebrate nociceptors^17^, we first sought to identify anoctamins which might selectively participate in *Drosophila* cold nociception.

Cell-type specific transcriptomic data indicate that the anoctamin genes *subdued* and *CG15270* are enriched in CIII cold nociceptors (**Table S1**; GEO accession GSE69353). In order to independently validate CIII expression of these genes, we performed qRT-PCR on isolated CIII neuron samples and found detectable expression of *subdued* (C_q_=24.53 ± 0.12) and *CG15270* (C_q_=28.29 ± 0.09). To further validate these data, we drove expression of GFP using *GAL4*s under control of the native *subdued* and *CG15270* promoters (**Figure 1A**), pairing these with a *Class III::tdTomato* fusion line. For *subdued*, we used the previously validated *GAL4*^*c240*^, which did in fact drive GFP expression in CIII nociceptors (**Figure 1B**). Consistent with previous reports^20^, *GAL4*^*c240*^ also drove GFP expression in CIV nociceptors (**Figure S1**). For *CG15270*, we made use of a *CG15270*-specific *T2A-GAL4* (Trojan *GAL4*); this approach has been shown to be a strong indicator of gene expression^34^. *CG15270 T2A-GAL4* drove expression of GFP in CIII nociceptors (**Figure 1C**). In fact, *CG15270* appeared to be broadly expressed in larval peripheral sensory neurons (**Figure S2A**), as well as in the larval central nervous system (**Figure S2B**).

**Figure 1.**
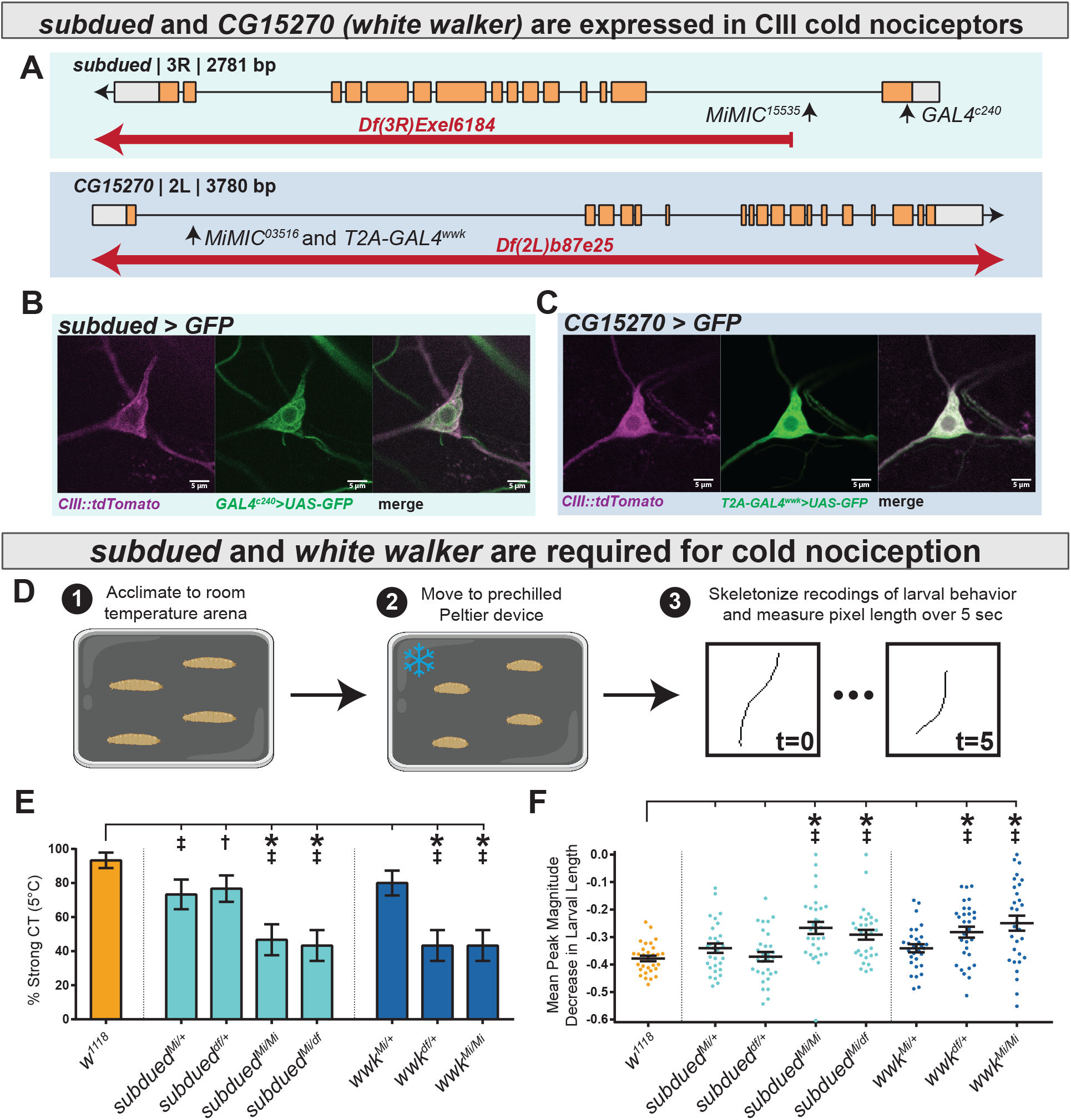
*subdued* and *white walker* (*CG15270*) are expressed in CIII neurons and required for cold nociception. (**A**) Alleles, chromosomal deficiencies, and gene-specific *GAL4s* used in this study. (**B-C**) *UAS-GFP* driven under the control of *GAL4*^*c240*^ and *T2A-GAL4*^*wwk*^ evidences that *subdued* and *white walker* are expressed in CIII neurons. (**D**) For cold plate assay, larvae were acclimated to a room temperature arena before being transferred to a pre-chilled cold plate. CT was identified by measuring the length of skeletonized larvae over the course of chilling. (**E**) % of animals which strongly CT in response to noxious cold (≥30% reduction in body length). Mutations in *subdued* and *white walker* result in a reduced percent of larvae which strongly CT in response to noxious cold (5°C). *w*^*1118*^ (n=30); *subdued*^*Mi/+*^ (n=30; p=0.13; BF_10_=4.417); *subdued*^*df/+*^ (n=30; p=0.25; BF_10_=2.826); *subdued*^*Mi/Mi*^ (n=30; p<0.001; BF_10_=461.34); *subdued*^*Mi/df*^ (n=30; p<0.001; BF_10_=997.24); *wwk*^*Mi/+*^ (n=30; p=0.45; BF_10_=2.00); *wwk*^*df/+*^ (n=30; p<0.001; BF_10_=997.24); *wwk*^*Mi/Mi*^ (n=30; p<0.001; BF_10_=997.24). (**F**) Mean peak magnitude in larval contraction, corresponding to panel E. *w*^*1118*^ (n=30); *subdued*^*Mi/+*^ (n=30; p=0.58; BF_10_=1.18); *subdued*^*df/+*^ (n=30; p=1; BF_10_=0.28); *subdued*^*Mi/Mi*^ (n=30; p<0.001; BF_10_=228.63); *subdued*^*Mi/df*^ (n=30; p=0.008; BF_10_=737.03); *wwk*^*Mi/+*^ (n=30; p=0.58; BF_10_=1.60); *wwk*^*df/+*^ (n=30; p=.003; BF_10_=324.11); *wwk*^*Mi/Mi*^ (n=30; p<0.001; BF_10_=433.18).

Using the previously developed cold plate assay^35^ we recorded larval behavior while delivering an acute ventral cold stimulus, which causes animals to contract (CT)^14^ along the head-to-tail axis (**Figure 1D**). Larvae carrying homozygous *Minos*-mediated integration cassette alleles^36^ (MiMIC, abbreviated as Mi) for *subdued* and *CG15270* (**Figure 1A**) showed decreased cold sensitivity – fewer animals strongly CT in response to noxious cold (**Figure 1E, Figure S3**) and homozygous larvae had a reduced peak magnitude of CT (measured by maximum magnitude reduction in length normalized to initial length, **Figure 1F**). These data suggest that *CG15270* and *subdued* are required for cold nociception. We therefore suggest the name *white walker* (*wwk*) for *CG15270*, as *white walker* mutant larvae are less sensitive to cold pain.

Larvae bearing a heterozygous *subdued* allele or a single chromosome with a deficiency covering the gene (*df*) displayed largely normal cold sensitivity (Bayesian analyses indicating evidence of a modest effect on % strong CT in the heterozygous condition), while the mutant allele paired with the deficiency recapitulated the relatively strong homozygous phenotype, thereby providing evidence against the phenotype being the result of off-target mutations.

Mutants carrying a heterozygous *white walker* allele behaved relatively normally, but a single deficiency resulted in reduced cold sensitivity. Further, the homozygous deficiency and the deficiency over MiMIC allele were lethal, meaning we could not rule out off-target effects by this approach alone.

### *subdued* and *white walker* are required in CIII neurons for CT behavior and cold-evoked neural activity

As *subdued* and *white walker* mutants had impaired cold nociception, we next sought to test the hypothesis that *subdued* and *white walker* function in CIII neurons (and further, to rule out off-target effects which could possibly be present in mutant backgrounds). To this end, we next knocked down the expression of *subdued* and *white walker* by *UAS*-mediated RNAi, targeting CIII neurons using the previously validated *GAL4*^*19-12*^ driver^14^. UP-TORR (https://www.flyrnai.org/up-torr/) was used to assess computationally predicted off-target effects for these RNAi constructs ^37^; UP-TORR indicates that these constructs have no predicted off-target effects. Consistent with the mutant phenotypes, CIII-specific knockdown of *subdued* or *white walker* resulted in reduced % CT (**Figure 2A, Figure S4**) and reduced peak magnitude of CT (**Figure 2B**). Although *subdued RNAi-2* showed a statistically insignificant decrease in % strong CT, the phenotype is extremely similar to *subdued RNAi-1* and Bayesian analyses indicate that there is substantial evidence in favor of the difference. For further experimentation we made use of the RNAi line with the strongest phenotype.

**Figure 2.**
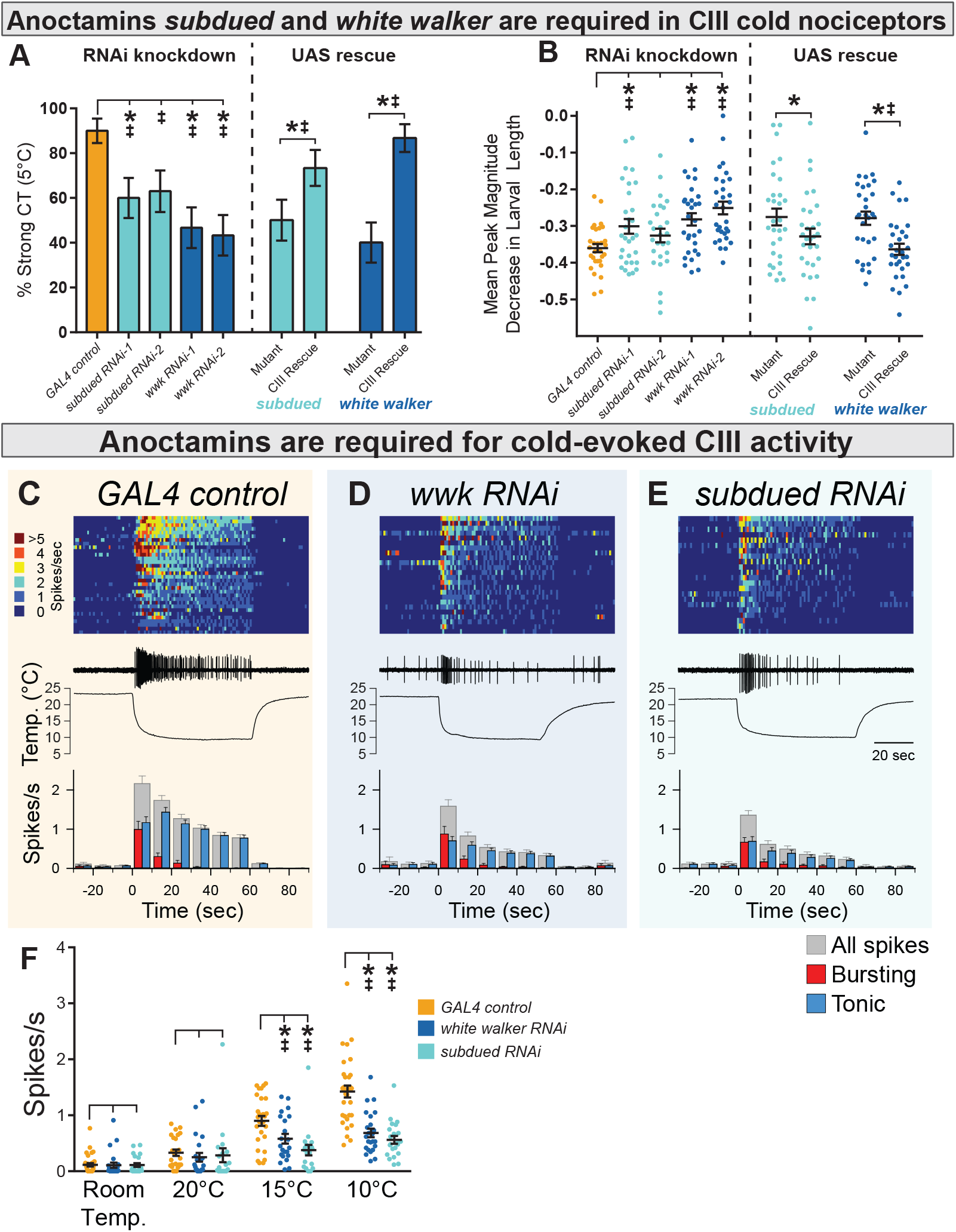
Anoctamins function in CIII neurons. (**A**) % of animals which strongly CT in response to noxious cold (≥30% reduction in body length). CIII-specific knockdown (*GAL4*^*19-12*^) of *subdued* and *white walker* result in a reduced percent of larvae which strongly CT in response to noxious cold. GAL4 control (n=30); *subdued RNAi-1* (n=30; p=0.039; BF_10_=7.64); *subdued RNAi-2* (n=27; p=0.075; BF_10_=4.93); *wwk RNAi-1* (n=30; p=.002; BF_10_=71.17); *wwk RNAi-2* (n=30; p<0.001; BF_10_=138.15). GAL4-UAS-mediated CIII rescue of *subdued* and *white walker* in mutant backgrounds increased cold sensitivity. *GAL4*^*nompC*^*;subdued*^*Mi/Mi*^ (n=30); *GAL4*^*nompC*^*/UAS-subdued;subdued*^*Mi/Mi*^ (n=30; p=0.031; BF_10_=3.57). *wwk*^*Mi/Mi*^*;GAL4*^*19-12*^ (n=30); *wwk*^*Mi/Mi*^*;GAL4*^*19-12*^*/UAS-wwk* (n=30; p<0.001; BF_10_=281.95). (**B**) Mean peak magnitude in larval contraction, corresponding to panel A. Knockdown: GAL4 control (n=30); *subdued RNAi-1* (n=30; p=0.049; BF_10_=3.98); *subdued RNAi-2* (n=27; p=0.437; BF_10_=0.78); *wwk RNAi-1* (n=30; p=.005; BF_10_=89.25); *wwk RNAi-2* (n=30; p<0.001; BF_10_=6932.18). Rescue: *GAL4*^*nompC*^*;subdued*^*Mi/Mi*^ (mutant, n=30); *GAL4*^*nompC*^*/UAS-subdued;subdued*^*Mi/Mi*^ (CIII Rescue, n=30; p=0.049; BF_10_=1.61). *wwk*^*Mi/Mi*^*;GAL4*^*19-12*^ (mutant, n=30); *wwk*^*Mi/Mi*^*;GAL4*^*19-12*^*/UAS-wwk* (CIII rescue, n=30; p<0.001; BF_10_=83.59). (**C, D, E**) Top: Heatmap representation of cold-evoked CIII activity (10°C), with each line representing an individual sample prep. Middle: Representative traces of cold-evoked neural activity over graph of temperature ramp. Knockdown of *subdued* and *white walker* results in decreased cold-evoked firing, particularly in the steady-state tonic firing at steady-state temperature. Bottom: Representation of average frequency from population binned by 10 sec. Red and blue bars show the proportion of bursting vs tonic spiking activity. Knockdown of *subdued* and *white walker* results in visibly decreased tonic firing. (**F**) CIII-specific knockdown of *subdued* and *white walker* results in decreased cold-evoked firing frequency at temperature ramps to 15°C and 10°C. Room Temp (N=78): GAL4 control (n=32); *white walker RNAi* (n=24; p>0.99; BF_10_=0.274); *subdued RNAi* (n=22; p=>0.99; BF_10_=0.279). 20°C (N=65): GAL4 control (n=25); *white walker RNAi* (n=22; p=0.69; BF_10_=0.39); *subdued RNAi* (n=18; p=0.90; BF_10_=0.32). 15°C (N=68): GAL4 control (n=27); *white walker RNAi* (n=22; p=0.0082; BF_10_=3.64); *subdued RNAi* (n=19; p<0.001; BF_10_=92.73). 10°C (N=78): GAL4 control (n=32); *white walker RNAi* (n=24; p<0.001; BF_10_=3310.09); *subdued RNAi* (n=22; p<0.001; BF_10_=44346.95).

We further tested the hypothesis that *subdued* and *white walker* function in CIII neurons by assessing whether the mutant phenotypes were due to loss of function in CIII nociceptors, predicting that CIII driven *UAS-white walker* or *UAS-subdued* expression would rescue CT in mutant backgrounds. For these experiments, we used CIII-targeting *GAL4*s located on chromosomes other than the chromosome containing the experimental gene (for *white walker, GAL4*^*19-12*^; for *subdued, GAL4*^*nompC*^). *GAL4-UAS*-mediated rescue in mutant backgrounds increased cold sensitivity (**Figure 2A-B, Figure S5**), indicating that *subdued* and *white walker* mutant phenotypes are due to selective anoctamin loss of function in cold nociceptors.

We next tested the hypothesis that anoctamins are involved in cold-evoked neural activity by making electrophysiological recordings in live, filet larvae during chilling. In control filet larvae, CIII firing frequency had an inverse relationship with temperature; chilling ramp resulted in bursting activity, while stable cold exposure resulted in tonic firing which increased in frequency with decreased temperature (**Figure 2C, 2F**). CIII-specific knockdown of *subdued* or *white walker* resulted in decreased cold-evoked CIII neural activity, particularly in steady-state tonic firing at a stable cold temperature (**Figure 2C-E**). Relative to controls, CIII-specific knockdown of *subdued* or *white walker* resulted in a reduction in overall firing frequency (spikes/sec) at both 15°C and 10°C (**Figure 2F**).

### *subdued* and *white walker* are not required for mechanosensation and do not play a role on dendritogenesis

We next tested the alternative hypothesis that these observed phenotypes reflect a loss of general excitability of CIII neurons. As CIII neurons are multimodal cold nociceptors and gentle touch mechanosensors, loss of general excitability or overall neural function would result in decreased gentle-touch mechanosensitivity; for example, previous reports have shown that loss of function of TRP channels required for cold nociception also results in defects in gentle-touch mechanosensitivity^14^. Innocuous touch assays (based on the Kernan assay^15^) revealed that neither *subdued* or *white walker* loss of function affects gentle-touch mechanosensitivity (**Figure 3A-B**). These results indicate that *subdued* and *white walker* are selectively required for CIII cold nociception.

**Figure 3.**
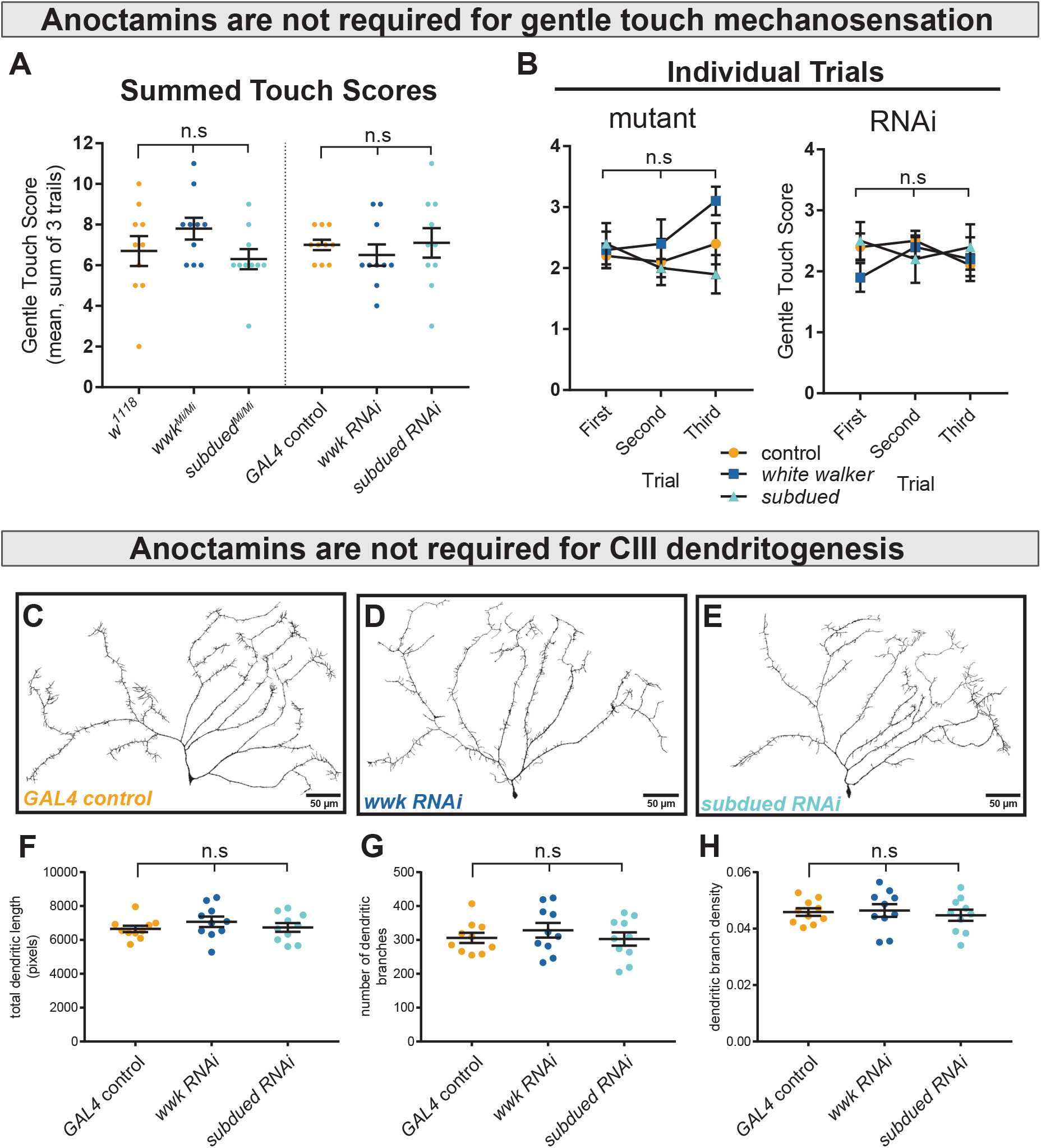
*subdued* and *white walker* are not required for innocuous mechanosensation or CIII dendritogenesis. (**A**) There is no difference in gentle touch mechanosensation (sum Kernan touch scores, 3 trials for each sample) in either *subdued* and *white walker* mutants (n=10 for each condition; p=0.20; BF_10_=0.62) or CIII-knockdown (n=10 for each condition; p=0.70; BF_10_=0.27). (**B**) For each genotype, there were no within-subjects effects in gentle touch sensation across trials. *w*^*1118*^ (n=10; p=0.69; BF_10_=0.27); *wwk*^*Mi/Mi*^ (n=10; p=0.19; BF_10_=0.85); *subdued*^*Mi/Mi*^ (n=10; p=0.42; BF_10_=0.42); *GAL4* control (n=10; p=0.39; BF_10_=0.50); *wwk RNAi* (n=10; p=0.37; BF_10_=0.47); *subdued RNAi* (n=10; p=0.80; BF_10_=0.25). (**C-E**) Representative neural traces of CIII ddaF neuron dendritic arbors under control and knockdown conditions. (**F**) Measures of total dendritic length show no differences between control and knockdown (n=10 for each condition; p=0.48; BF_10_=0.34). (**G**) There is no difference in the number of branches between control and knockdown (n=10 for each condition; p=0.70; BF_10_=0.27). (**H**) Dendritic branch density does not differ between control and knockdown (n=10 for each condition; p=0.81; BF_10_=0.24).

**Figure 4.**
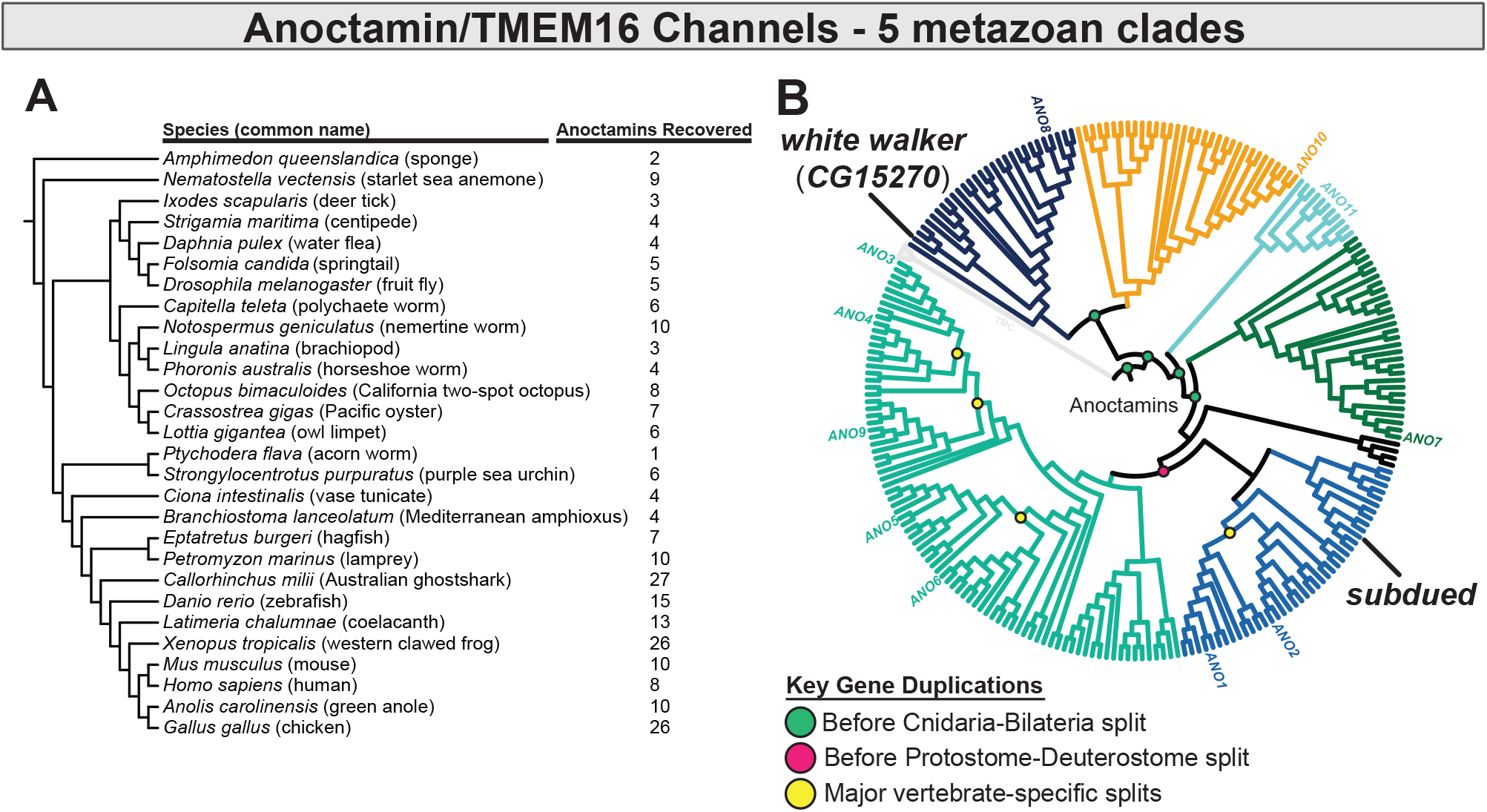
Anoctamin/TMEM16 channels form 5 metazoan and 6 nephrozoan clades. (**A**) Species, cladogram, and number of sequences used in this analysis. (**B**) Maximum likelihood phylogeny of animal anoctamins, rectified and rearranged against cladogram in A. *subdued* is a member of the ANO1/2 clade of nephrozoan calcium-activated chloride channels, and of the clade of metazoan anoctamins that includes mammalian ANO1/2/3/4/5/6/9. Mammalian ANO8 is homologous to *white walker*. Additionally, this phylogeny evidences a separate clade of ANOs of unknown function we have deemed ANO11 (cyan).

We also tested the alternative hypothesis that decreased cold sensitivity was due to morphological defects in CIII dendritic arborization. CIII neurons labeled with GFP show no dendritic defects under *subdued* and *white walker* knockdown (**Figure 3C-E**); there were no quantitative differences in the number of dendritic branches, total dendritic length, or dendritic branch density (**Figure 3F-H**).

### The evolutionary history of anoctamins supports that *subdued* is part of the ANO1/ANO2 subfamily a Ca^2+^-activated Cl^-^ channels

While previous work has phylogenetically characterized anoctamins/TMEM16 channels ^38-40^, their evolution and familial organization outside of vertebrates is relatively poorly understood, and therefore limits our ability to formulate hypotheses of function based on phylogeny. We therefore generated an anoctamin phylogeny which is inclusive of a wide range of animal taxa (**Figure 4; Figures S6-S7**). This phylogeny indicates that anoctamins are organized into 5 major metazoan subfamilies which predate the Cnidaria-Bilateria split (including a new subfamily we deem ANO11, which has not been previously described), and 6 major bilaterian subfamilies. *subdued* has been previously determined to encode a Ca^2+^- activated Cl^-^ channel (CaCC) ^19^; consistent with these findings, this phylogeny evidences that *subdued* is a member of the bilaterian ANO1/2 subfamily of CaCCs. *white walker* is one of the more distantly related anoctamins and is of unknown function, but in contrast to *subdued*, has a single homologue in humans (both part of the metazoan ANO8 subfamily).

### CIII neurons make use of excitatory Cl^-^ currents to encode noxious cold

As *subdued* encodes an ANO1/ANO2 orthologue and Ca^2+^-activated Cl^-^ channel^19^, and positively regulates cold nociception, we hypothesized that CIII neurons make use of atypical, excitatory Cl^-^ currents to encode noxious cold.

Across animal taxa, neural Cl^-^ homeostasis is maintained by differential expression of SLC12 co-transporters – in *Drosophila, kazachoc* (*kcc*) encodes an outwardly facing K^+^-Cl^-^ cotransporter ^41^, while *ncc69* encodes an inwardly facing Na^+^-K^+^-Cl^−^ cotransporter ^42^. The differential expression of these cotransporters can therefore modulate the membrane-potential effects of Cl^-^ currents. Transcriptomic data indicates that *kcc* is downregulated and *ncc69* is upregulated in CIII neurons (**Table S1**), indicating that CIII neurons may maintain relatively high intracellular Cl^-^ concentrations, thereby facilitating depolarizing (and therefore excitatory) Cl^-^ currents. In order to manipulate Cl^-^ homeostasis in CIII nociceptors we used RNAi and UAS constructs to knock down and overexpress *ncc69* and *kcc*, and thereafter observed effects on noxious cold-evoked behavior.

Modulating the expression of SLC12 cotransporters affected cold-evoked behavior (**Figure 5A-B, Figure S8**). Knockdown of *ncc69* and overexpression of *kcc—*both of which would theoretically decrease intracellular Cl^-^ concentration*—*resulted in reduced % strong CT. This difference was statistically different for *ncc69 RNAi*, and there is substantial evidence according to Bayesian analyses for an effect in both *ncc69 RNAi* and *kcc OE* on % strong CT. However, support for the *kcc OE* phenotype was notably weaker, as there was no evidence for differences in average peak magnitude decrease in larval length. Further, knockdown of *kcc* and overexpression of *ncc69—*which would both theoretically increase intracellular Cl^-^ concentration*—*did not obviously affect sensitivity to 5°C noxious cold. Manipulating Cl^-^ physiology in this fashion did not affect gentle touch mechanosensitivity, indicating that Cl^-^ physiology is selectively required for CIII cold nociception (**Figure S9**)

**Figure 5.**
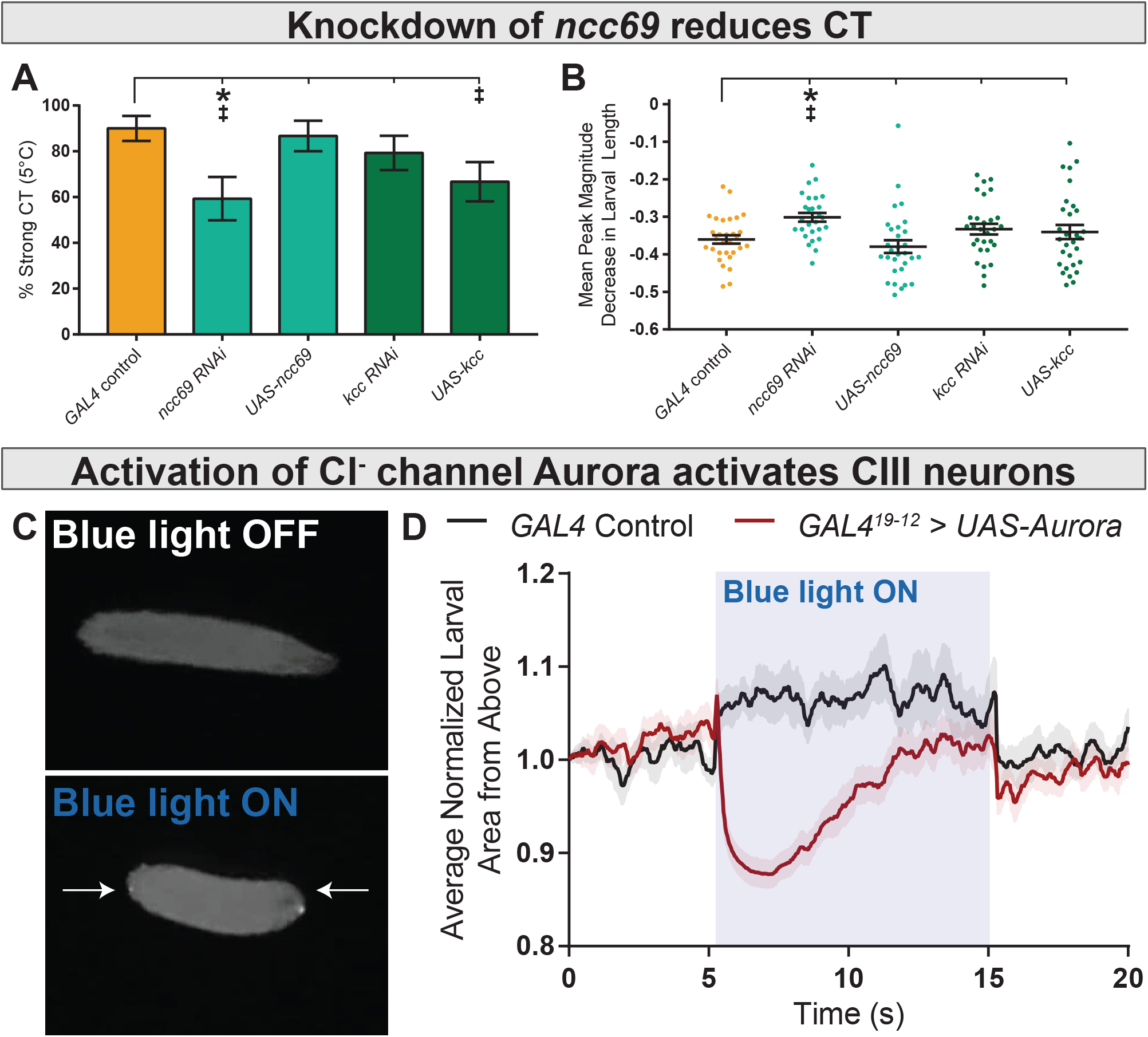
CIII Cl^*-*^ currents are excitatory. (**A**) % of animals which strongly CT in response to noxious cold (≥30% reduction in body length). CIII-specific knockdown (GAL4^19-12^) of *ncc69* results in a reduced percent of larvae which strongly CT in response to noxious cold. GAL4 control (n=30); *ncc69 RNAi* (n=29; p=0.038; BF_10_=7.96); *ncc69 OE* (n=30; p=1; BF_10_=0.58); *kcc RNAi* (n=29; p=0.91; BF_10_=0.99); *kcc OE* (n=30; p=0.13; BF_10_=3.29). (**B**) Mean peak magnitude in larval contraction, corresponding to panel A. GAL4 control (n=30); *ncc69 RNAi* (n=29; p=0.025; BF_10_=46.38); *ncc69 OE* (n=30; p=0.77; BF_10_=0.38); *kcc RNAi* (n=29; p=0.51; BF_10_=0.68); *kcc OE* (n=30; p=0.76; BF_10_=0.37). (**C**) Optogenetic activation of CIII>Aurora activates CIII neurons, resulting in CT behavior. (**D**) CT behavior for control and CIII>Aurora represented as larval area from above. Blue light activation of the Cl^-^ channel Aurora causes a rapid reduction in normalized larval area from above, an indication of CT behavior. Comparison of minimum larval area measured during blue light illumination: *GAL4* control (n=10); *GAL4*^*19-12*^ *> UAS-Aurora* (n=20; p<0.001; BF_10_=134.78).

These results are consistent with the hypothesis that CIII Cl^-^ currents are excitatory and selectively required for cold. However, these results are not alone sufficient to disprove the alternative hypothesis that Cl^-^ currents are inhibitory; for example, inhibitory currents may be required for patterning neural activity in such a way to appropriately drive behavior, as is the case in CIV neurons with respect to hyperpolarizing K^+^ currents ^43,44^.

In order to further test these competing hypotheses, we drove CIII expression of Aurora, an engineered Cl^-^ channel which gates in response to blue-light illumination ^45^. Typically, Aurora is employed as an inhibitory optogenetic tool. If CIII Cl^-^ currents are excitatory, one would expect optogenetic activation of these Cl^-^ channels to result in CIII-mediated behaviors. In fact, blue-light illumination of freely locomoting *GAL4*^*19-12*^*>UAS-Aurora* larvae elicited CT behavior (frequently followed by head-casting and reverse locomotor behavior), indicating that Cl^-^ currents are excitatory and activate CIII neurons (**Figure 5C-D**). These findings collectively disprove the hypothesis that cold-evoked CIII Cl^-^ currents are inhibitory.

### Low extracellular Cl-concentrations and overexpression of *ncc69* results in nociceptive sensitization

We next modulated extracellular Cl^-^ in our electrophysiology preparations in order to more directly observe the effect of the Cl^-^ gradient on CIII activity. Decreasing extracellular Cl^-^ sensitized CIII neurons; bath application of low Cl^-^ saline induced spontaneous CIII bursting and increased sensitivity to cooling (**Figure 6**), effects which could be washed out. Relative to controls, we observed increased spontaneous firing under low Cl^-^ saline at room temperature, increased firing at 20°C and 15°C, and no effect at 10°C (**Figure 6B**).

**Figure 6.**
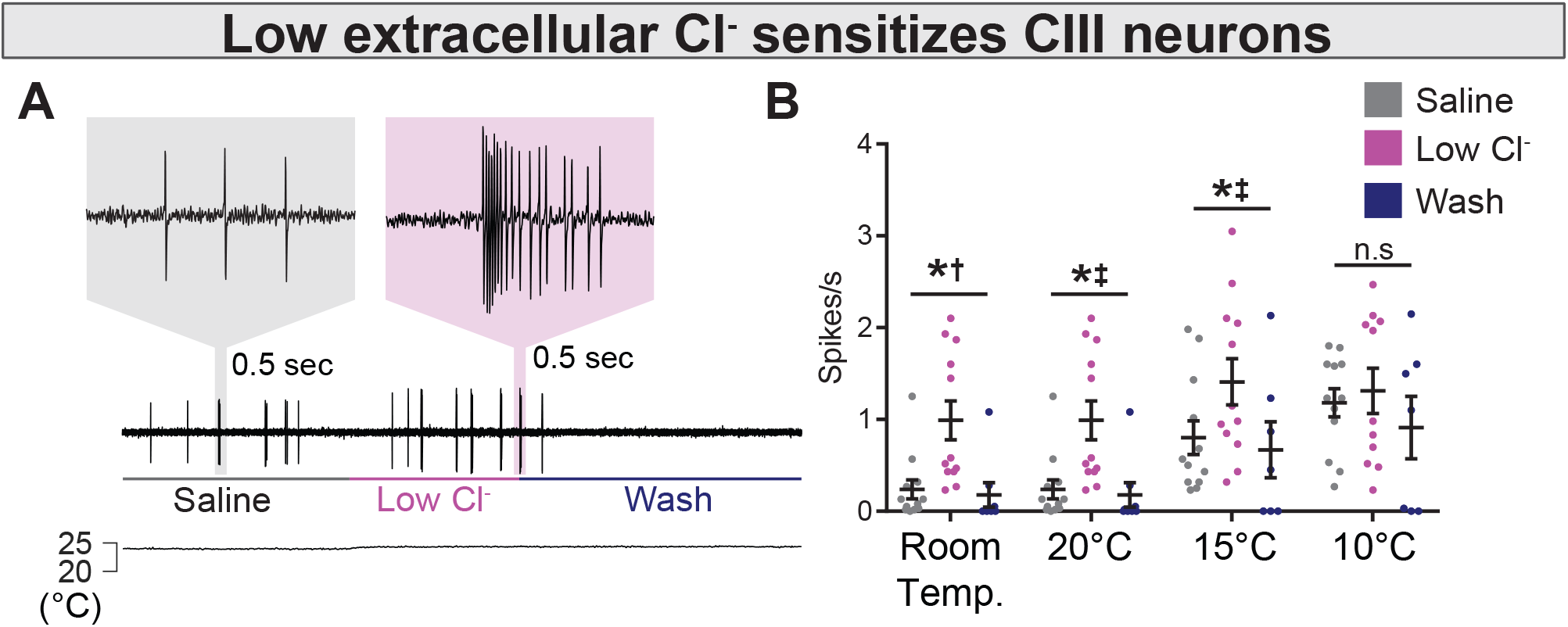
Modulating extracellular Cl^-^ sensitizes CIII neurons. (**A**) Extracellular application of low Cl^-^ saline to fileted electrophysiology preps causes spontaneous bursting activity. At room temperature, neurons are largely silent, but may show occasional, low frequency spiking (likely associated with mechanosensation from saline flow). (**B**) The presence of low Cl^-^ saline causes spontaneous bursting and sensitizes CIII neurons to cooling. Room Temp: saline (n=12); low Cl^-^ (n=12); wash (n=8); p=0.003, BF_10_=1.89. 20°C: saline (n=12); low Cl^-^ (n=12); wash (n=8); p=0.003, BF_10_=66.38. 15°C: saline (n=12); low Cl^-^ (n=12); wash (n=7); p=0.011, BF_10_=14.51. 10°C: saline (n=12); low Cl^-^ (n=11); wash (n=7); p=0.25, BF_10_=0.75.

One of the hallmarks of neuropathic pain following spinal cord injury in mammals is upregulation of *NKCC1* (a human orthologue of *ncc69*) in neurons involved in nociception, resulting in increased intracellular Cl^-^, and thereby aberrantly excitatory Cl^-^ currents, leading to nociceptive sensitization and spontaneous nociceptor activity, much like we observed under low Cl^-^ saline conditions ^46,47^. In an attempt to genetically model neuropathic pain we overexpressed *ncc69* in CIII neurons, predicting it would likewise cause nociceptive hypersensitivity.

Overexpression of *ncc69* sensitized neurons to cooling and resulted in spontaneous, room temperature CIII activity (**Figure 7A, 7C, 7D**). Curiously, we did not observe deficits in CIII firing under *ncc69* knockdown (**Figure 7B**).

**Figure 7.**
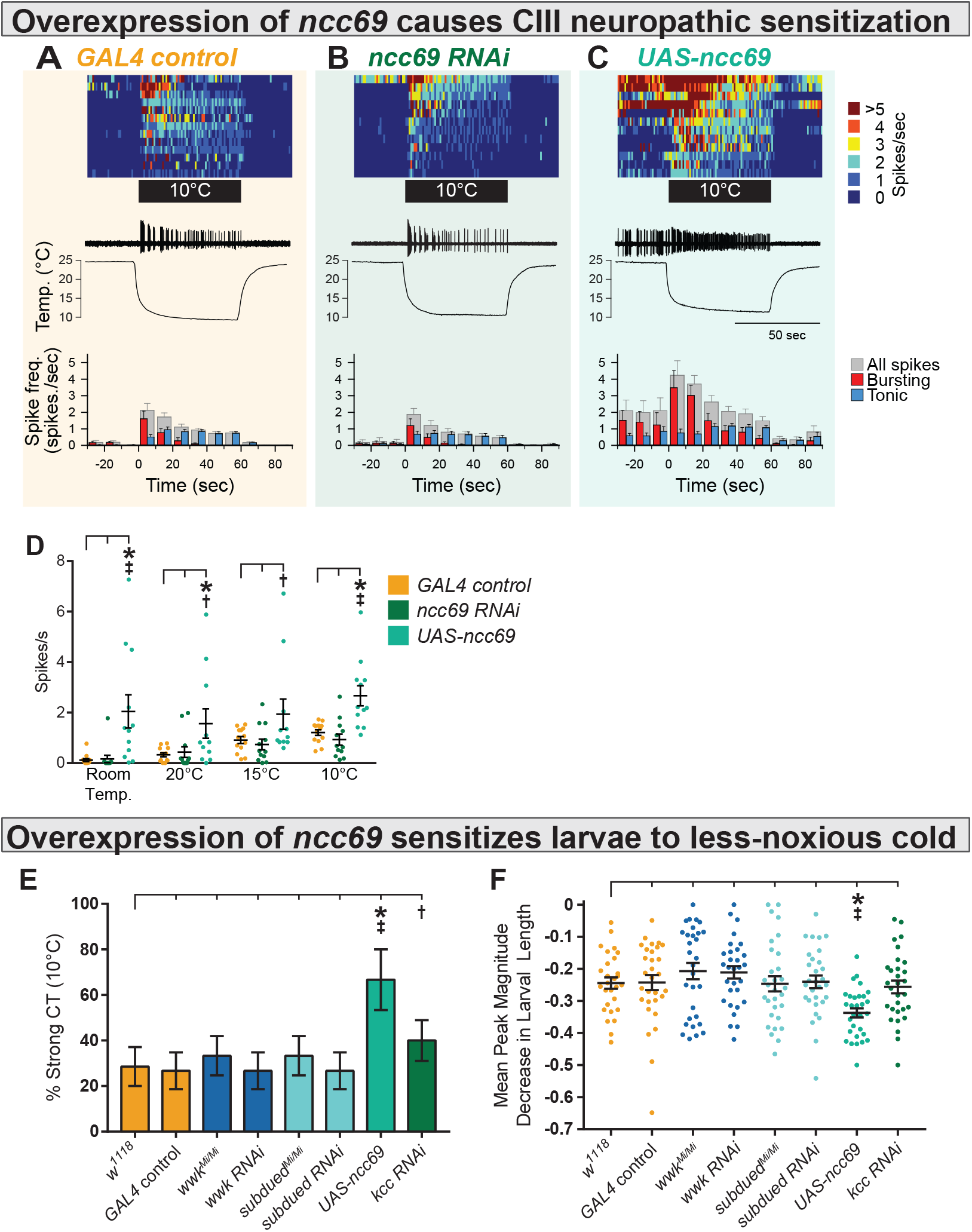
Overexpression of *ncc69* causes nociceptive hypersensitization. (**A, B, C**) Top: Heatmap representation of cold-evoked CIII activity, with each line representing an individual sample prep. Middle: Representative traces of cold-evoked neural activity over graph of temperature ramp. Overexpression of ncc69 results in spontaneous nociceptor activity and increased cold sensitivity. Bottom: Representation of average spike frequency from population binned by 10 sec. Red and blue bars show the proportion of bursting vs tonic spiking activity. (**D**) Overexpression of *ncc69* causes spontaneous neural activity and sensitizes neurons to cooling. Room Temp: *GAL4* control (n=13); *ncc69* RNAi (n=12; p=0.99; BF_10_=0.22); *UAS-ncc69* (n=12; p<0.001; BF_10_=7.86). 20°C: *GAL4* control (n=13); *ncc69* RNAi (n=12; p=0.96; BF_10_=0.40); *UAS-ncc69* (n=11; p=0.020; BF_10_=2.23). 15°C: *GAL4* control (n=13); *ncc69* RNAi (n=12; p=0.90; BF_10_=0.26); *UAS-ncc69* (n=11; p=0.062; BF_10_=1.17). 10°C: *GAL4* control (n=13); *ncc69* RNAi (n=12; p=0.77; BF_10_=0.60); *UAS-ncc69* (n=11; p=0.0042; BF_10_=25.18). (**E**) % of animals which strongly CT in response to noxious cold (10°C). CIII-specific (*GAL4*^*19-12*^) overexpression of *ncc69* results in an increased percentage of strong CT in response to less-noxious cold. *w*^*1118*^ (n=28); *GAL4 control* (n=30; p=1; BF_10_=0.46); *wwk*^*Mi/Mi*^ (n=30; p=1; BF_10_=0.69); *wwk RNAi* (n=30; p=1; BF_10_=0.46); *subdued*^*Mi/Mi*^ (n=30; p=1; BF_10_=0.69); *subdued RNAi* (n=30; p=1; BF_10_=0.46); *UAS-ncc69* (n=30; p=0.026; BF_10_=28.79); *kcc RNAi* (n=30; p=1; BF_10_=1.07). (**F**) Mean peak magnitude in larval contraction, corresponding to panel E. *w*^*1118*^ (n=28); *GAL4 control* (n=30; p=1; BF_10_=0.27); *wwk*^*Mi/Mi*^ (n=30; p=1; BF_10_=0.48); *wwk RNAi* (n=30; p=1; BF_10_=0.52); *subdued*^*Mi/Mi*^ (n=30; p=1; BF_10_=0.27); *subdued RNAi* (n=30; p=1; BF_10_=0.27); *UAS-ncc69* (n=30; p=0.037; BF_10_=199.79); *kcc RNAi* (n=30; p=1; BF_10_=0.29).

We next assessed cold nociception in response to less-noxious cold (10°C), which causes strong CT in only ∼30% of wild-type animals and is not affected by *subdued* or *white walker* loss of function (**Figure 7E-F, Figure S10**). Based on our electrophysiological observations, we predicted that overexpression of *ncc69* and knockdown of *kcc* would result in cold hypersensitivity to less-noxious temperature. Consistent with these electrophysiological observations, *GAL4-UAS*-mediated overexpression of *ncc69* resulted in increased cold sensitivity at less-noxious temperatures; more subjects strongly CT in response to less-noxious cold (**Figure 7E, Figure S10**) and the magnitude of CT was substantially stronger (**Figure 7F**). However, larvae were not sensitized to innocuous touch (**Figure S9**), and unexpectedly, the cold-sensitivity phenotype was not mirrored by *kcc* knockdown (a sensitization phenotype only barely more likely than no phenotype, as inferred by Bayesian analyses).

## DISCUSSION

Here, we have shown that CIII cold nociceptors make use of excitatory Cl^-^ currents in order to selectively encode cold. Our current working hypothesis in light of these findings is that cold-evoked, TRP-channel mediated Ca^2+^ currents activate Ca^2+^-activated Cl^-^ channels (CaCCs), which due to differential expression of *ncc69* and *kcc*, results in depolarizing Cl^-^ currents, enhancing neural activation in response to cold (**Figure 8**). These results support a role for *subdued, white walker*, and *ncc69* in selectively facilitating CIII-dependent cold nociception and not mechanosensation, thereby participating in mechanisms that allow CIII neurons to differentiate between sensory modalities.

**Figure 8.**
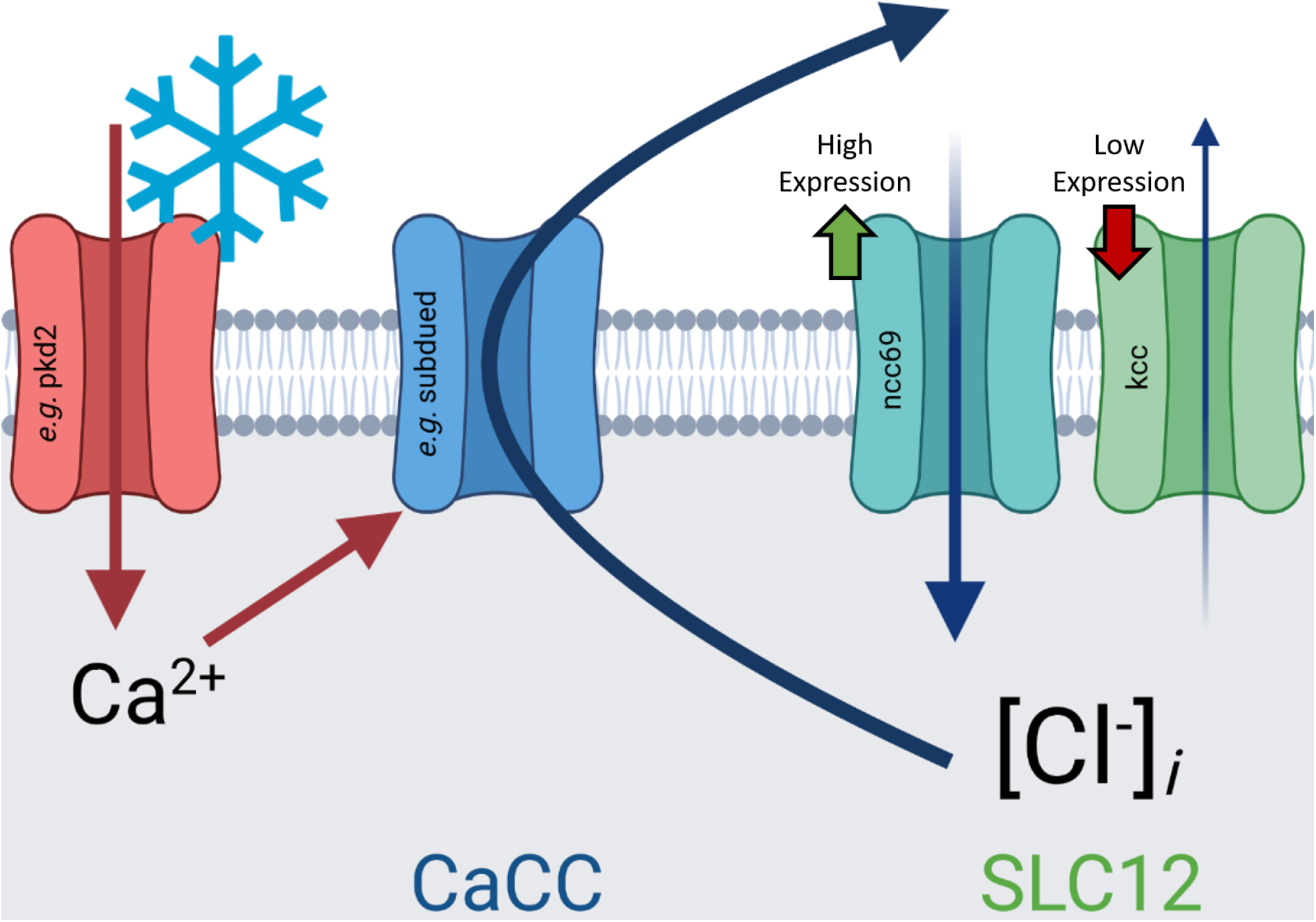
Graphical summary of hypothesis outlined in discussion: cold-evoked, TRP-channel mediated Ca^2+^ currents activate Ca^2+^-activated Cl^-^ channels (CaCCs), which due to differential expression of *ncc69* and *kcc*, results in depolarizing Cl^-^ currents, enhancing neural activation in response to cold.

As *subdued* has been previously characterized as a CaCC, its role is consistent with the hypothesis outlined above. However, the evolution of *subdued* has been implicitly debated in the literature ^18,19^, with suggestion that it may be more closely related to ANO6 ^18^. Our phylogenetic analysis strongly evidences that *subdued* is part of the bilaterian ANO1/ANO2 subfamily of CaCCs. Moreover, our phylogeny suggests that insects have no direct ANO6 homologue, as the diversification of ANO3, ANO4, ANO5, ANO6, and ANO9 occurred after the protostome-deuterostome split. The role of *subdued* in cold nociception therefore may constitute functional homology in the bilaterian ANO1/ANO2 subfamily, as mammalian ANO1 has been shown to participate in nociception alongside mammalian TRP channels ^48^. However, the possibility of convergent evolution cannot yet be ruled due to the absence of evidence of function in other taxa.

In contrast, *white walker* has not been demonstrated to function as, or be closely related to, CaCCs. Our phylogeny evidences that *white walker* is part of the metazoan ANO8 subfamily; one important function of mammalian ANO8 is to tether the endoplasmic reticulum (ER) and plasma membrane (PM), thereby facilitating inter-membrane Ca^2+^ signaling ^49^. Therefore, a speculative hypothesis is that *white walker* likewise serves to couple the ER and PM, and that subsequently, ER-dependent Ca^2+^ signaling might promote the CIII cold response. However, ANO8 has been shown to conduct Cl^-^ heterologously ^50^, so Cl^-^ channel function in *Drosophila* cannot be ruled out *a priori*. As *white walker* appears to be broadly expressed in neural tissues, *white walker* may function as a fundamental component of insect neural machinery, and is therefore likely to be a gene of interest in future studies.

In addition to the functions outlined above, anoctamins—including subdued^18^—are known to also function as lipid scramblases ^51^. A plausible alternative hypothesis is therefore that *subdued* and/or *white walker* function as lipid scramblases as part of unidentified signaling cascades critical to noxious cold transduction.

The results of our *ncc69* knockdown behavior, Cl^-^-channel optogenetics, and Cl^-^ electrophysiology experiments are consistent with the hypothesis that CIII neurons make use of atypical excitatory Cl^-^ currents. However, we did not observe an effect on cold-evoked CIII activity in response to *ncc69* knockdown (**Figure 7B**). As *ncc69* knockdown behavioral defects are seen at 5°C (**Figure 5A-B**) and not at 10°C (**Figure 7E-F**), it may be the case that this knockdown only affects electrical activity at very noxious temperatures. Our inability to detect deficiencies in cold-evoked neural activity may therefore be due to limitations in our electrophysiology prep, which limit our ability to cool below 10°C. These results are still curious, however, as *subdued* and *white walker* knockdowns result in electrophysiological defects at less-noxious (10°C) and innocuous (15°C) temperature drops (**Figure 2D-F**). Moreover, although we have substantial Bayesian evidence of an effect on % strong CT under *kcc* overexpression (**Figure 5A**), this difference was not evidenced by traditional frequentist statistics, the phenotype did not clearly mimic *ncc69* knockdown (**Figure 5A**), nor did we see a difference in the mean peak magnitude of CT response (**Figure 5B**). In totality, these results may suggest that CIII Cl^-^ homeostasis involves other cotransporters which can adapt in either function or expression in response to loss or gain of function. Given the importance of this system to behavior selection, CIII Cl^-^ homeostasis will make an interesting target for future experimentation.

Importantly, we have shown that overexpression of *ncc69*—a fly orthologue of *NKCC1*— is sufficient for driving a neuropathic pain-like state in larvae. As misexpression of *NKCC1* is a major factor associated with neuropathic pain in humans, we posit that altered larval cold nociception constitutes a new system in which to study neuropathic pain. Importantly, this system is wholly genetic, and does not require injury or other methods of invoking nociceptive hypersensitization, making it a high-throughput and easily accessible tool. Interestingly, RNAi knockdown of *kcc* did not mirror the *ncc69* overexpression phenotype. We speculate that this is because native *kcc* expression levels are low enough that knockdown does not sufficiently disrupt Cl^-^ homeostasis. This might also be because of hypothetical unknown mechanisms of compensation, as discussed above.

While it has been often stated that neuropathic pain is maladaptive, there is growing support for the hypothesis that neuropathic pain has its mechanistic bases in adaptive nociceptive hypersensitization – a mechanism by which organisms are more readily able to respond to danger following insult ^48,52-63^. Nerve injury has been previously shown to cause nociceptive hypersensitization in adult *Drosophila*, and has been hypothesized to be protective^57^. Moreover, it has been recently shown that nociceptive hypersensitization of CIII neurons is coincident with cold acclimation, the mechanism by which insects adapt to dips in temperature^64^. One speculative hypothesis is that changes in expression levels of SLC12 transporters underlie these shifts in cold acclimation-induced cold sensitivity. This would be consistent with a study demonstrating that a number of genes in *Drosophila* involved in ion homeostasis are differentially regulated following cold acclimation ^65^. If this speculation is veridical, insect thermal acclimation may serve as an example of how “maladaptive” injury and neuropathic sensitization can confer an adaptive advantage. It is therefore possible that these findings, and continued study, will lead to not only advances relevant to human health, but also better our understanding of nervous system evolution and the evolution of mechanisms underlying neuropathic sensitization and pain.

## Acknowledgments

This work is supported by NIH R01 NS115209-01 (to DNC and GSC). NJH was supported by NIH F31 NS117087-01, a GSU Brains & Behavior Fellowship, and a Kenneth W. and Georganne F. Honeycutt Fellowship. JML was supported by a GSU Brains & Behavior Fellowship. We thank Dr. Don van Meyel (McGill University), Dr. Mark Tanoye (UC Berkeley), Dr. Changsoo Kim (Chonnam National University), Dr. Chun Han (Cornell University), and the FlyLight Project (Janelia Research Campus) for providing *Drosophila* stocks.

## Author Contributions

Conceptualization, NJH and DNC; Methodology, NJH, AS, and DNC; Cold-plate assays, NJH, MNB, and TRG; Innocuous touch assays, NJH; Cell isolation at qRT-PCR, SB; Optogenetics, NJH and JML; Constructed custom optogenetics rig, AAP; Microscopy, NJH, JML, and AAP; Neural reconstructions and morphometric analyses, JML; Electrophysiology, AS; Phylogenetics, NJH; Statistics and other formal analyses, NJH and AS; Characterized *GAL4*^*CIII*^ and developed *CIII::tdTomato* fusion line, AAP; Developed *UAS-wwk*, NJH and SB; Writing – Original Draft, NJH; Writing – Review & Editing – NJH, AS, JML, AAP, MNB, SB, TRG, GSC, and DNC; Visualization, NJH, AS, JML, and GSC; Supervision, DNC; Funding acquisition, NJH, GSC, and DNC.

