## Supplemental Files for "Chloride-dependent mechanisms of multimodal sensory discrimination and neuropathic sensitization in *Drosophila*"

**Table S1.** Class III expression of *subdued, white walker, ncc69, and kcc* from cell-type specific microarray (GSE69353)[1], expressed as mean log fold-change difference between isolated CIII and whole-larval samples. Positive values indicate enrichment/upregulation and negative values downregulation.

| **Family** | **Transcript** | **CIII:Whole Larva (mean log FC)** |
|---|---|---|
| Anoctamin | *subdued* | 4.56 |
| | *white walker (CG15270)* | 1.41 |
| SLC12 | *ncc69* | 6.71 |
| | *kcc* | -1.37 |

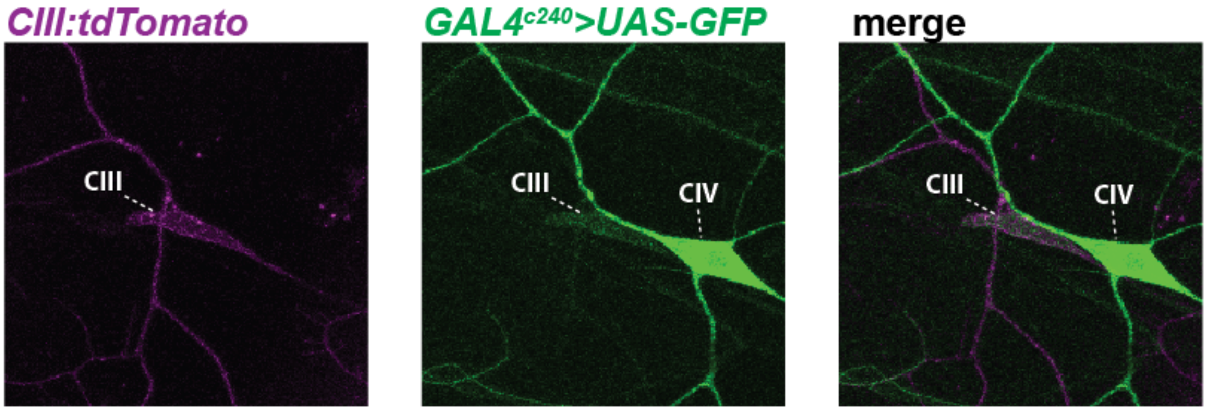


**Figure S1.** *GAL4^c240^* driven *UAS-GFP* expression. Consistent with previous reports[2], *GAL4^c240^* also shows strong expression in CIV neurons.

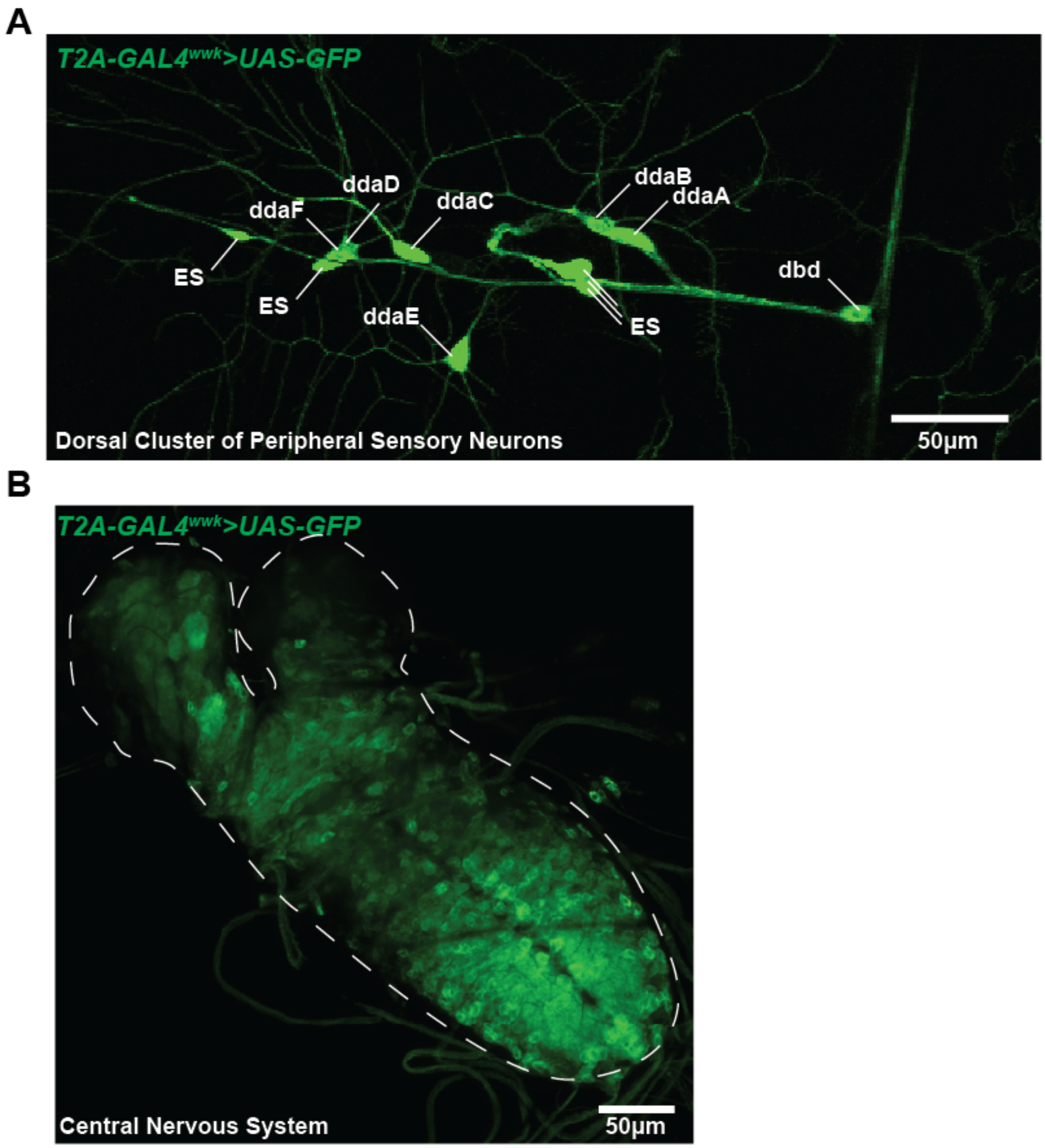


**Figure S2.** *T2A-GAL4^wwk* driven *UAS-GFP* expression. *T2A-GAL4^wwk* does not singularly drive expression in CIII neurons, but rather appears to broadly mark neural tissues. Examples: (**A**) *T2A-GAL4^wwk>GFP* expression patterns in the dorsal sensory neuron cluster. (**B**) *T2A-GAL4^wwk>GFP* in the larval central nervous system; expression appears through the central nerve cord and in restricted regions of the brain lobes.

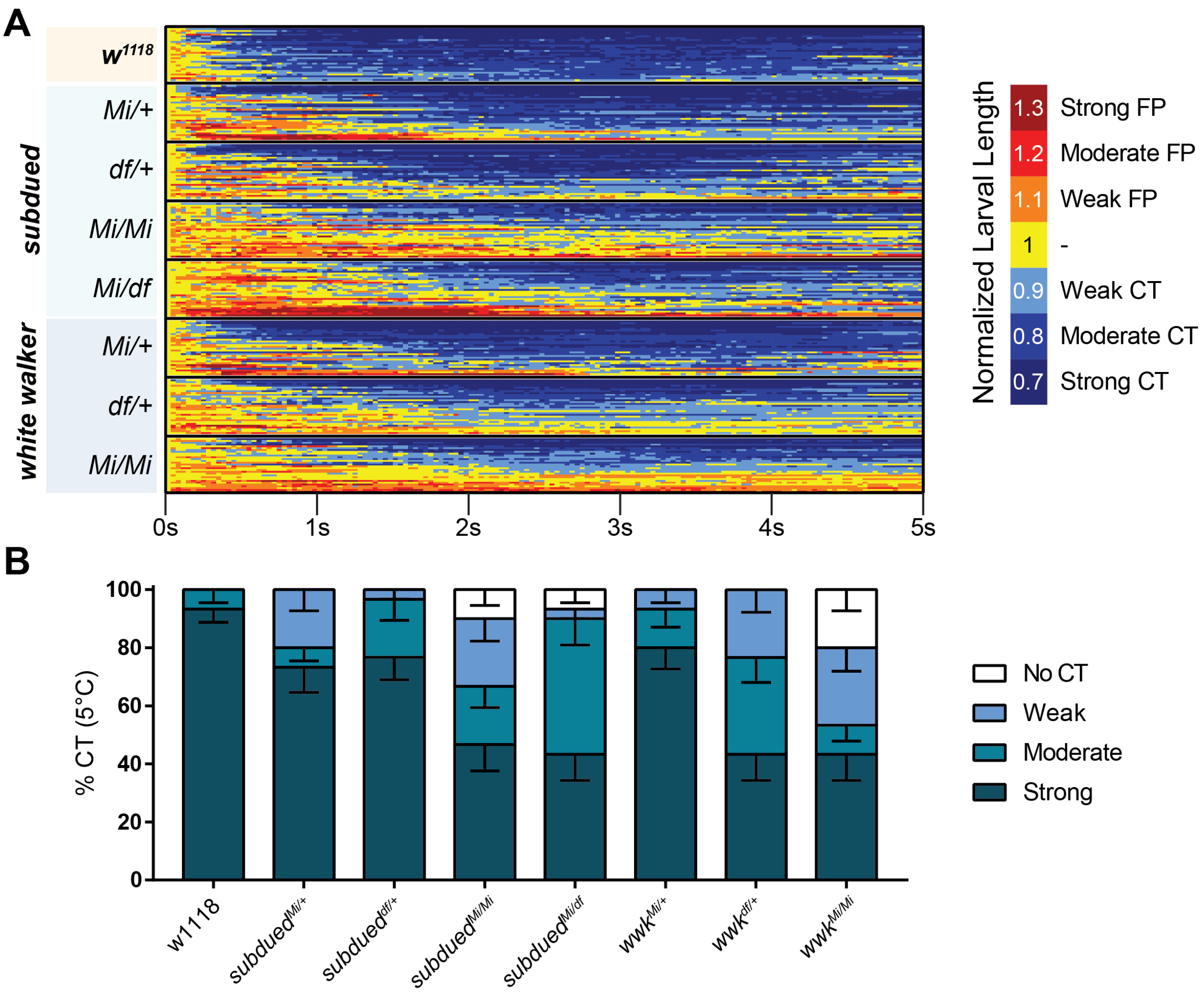


**Figure S3.** (**A**) Heatmap representation of cold-evoked CT in genetic control (*w*$^{1118}$) and mutant conditions, where FP is flaccid paralysis (full body relaxation with an inability to locomote) and CT is contraction. N=240, n=30 for each condition. (**B**) % of animals from anoctamin allele/deficiency experiments performing a particular CT response, where NR is No Response (<10% reduction in length), Weak is ≥10% reduction in length, Moderate is ≥20% reduction in length, and Strong is ≥30% reduction in length; bars show proportions in % ± SEP.

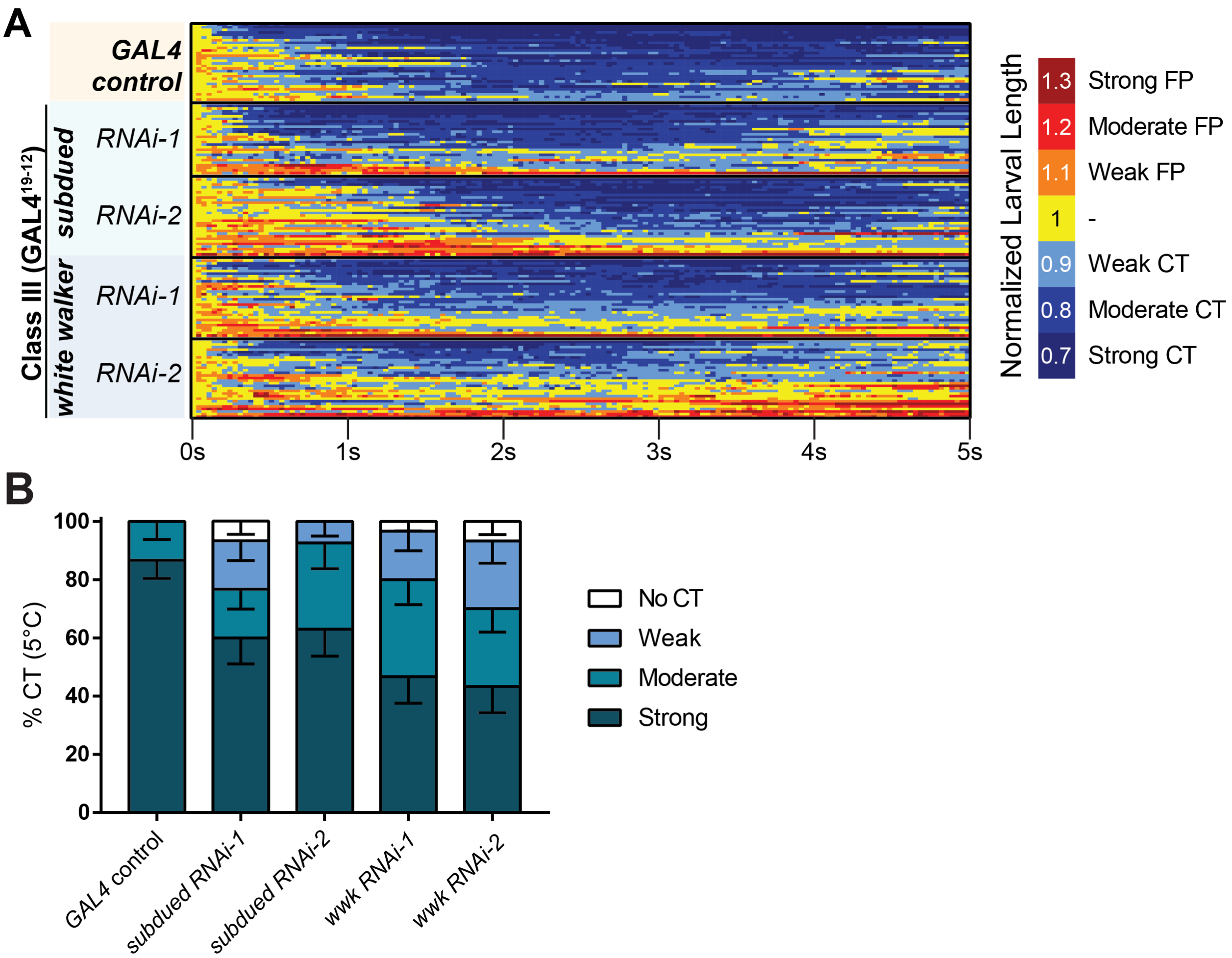


**Figure S4.** (**A**) Heatmap representation of cold-evoked CT in genetic control (*GAL4*) and knockdown conditions, where FP is flaccid paralysis and CT is contraction. N=147, n=30 for each condition except *subdued RNAi-2*, where n=27. (**B**) % of animals from anoctamin knockdown experiments performing a particular CT response, where NR is No Response (<10% reduction in length), Weak is ≥10% reduction in length, Moderate is ≥20% reduction in length, and Strong is ≥30% reduction in length; bars show proportions in % ± SEP.

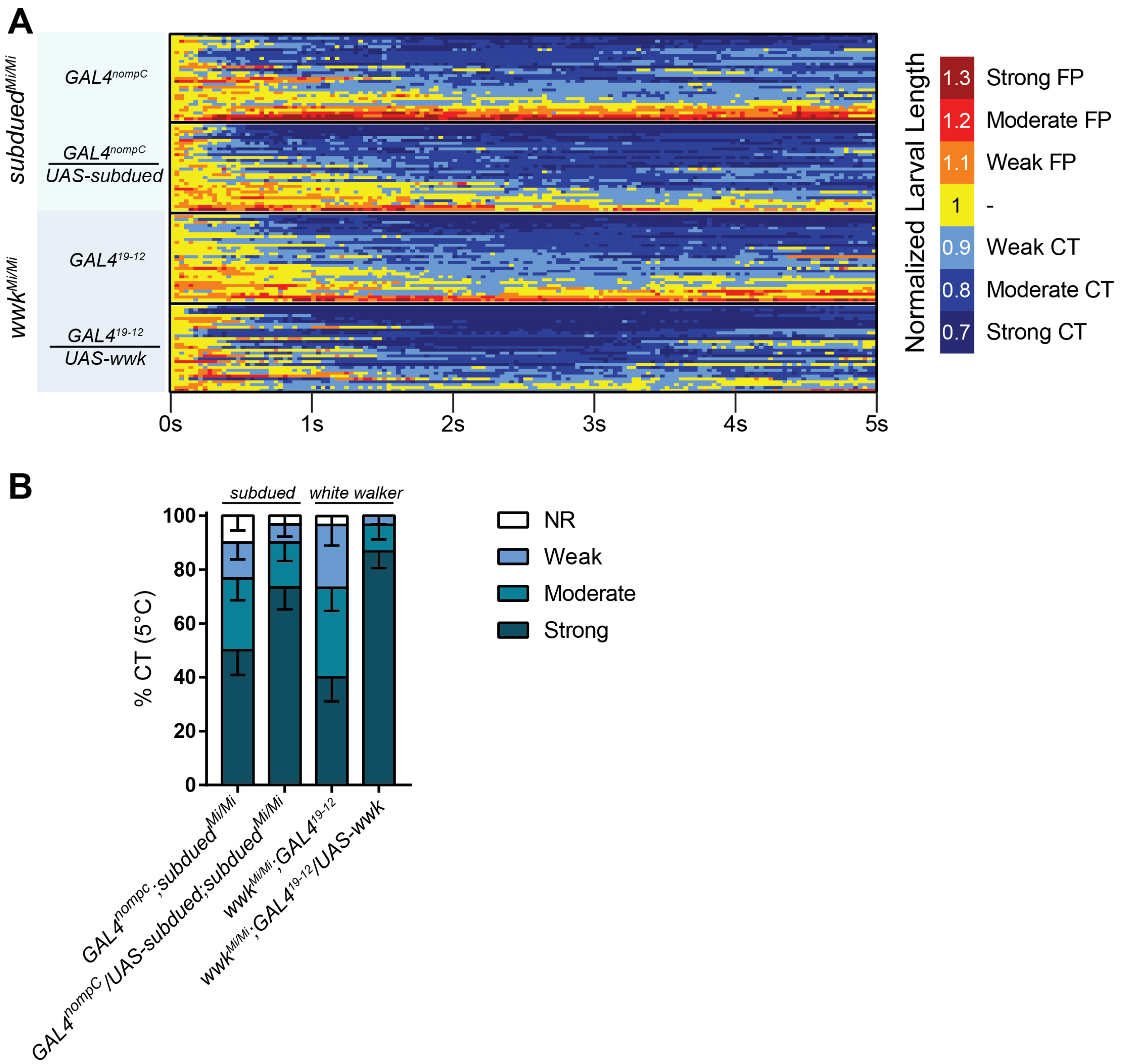


**Figure S5.** (**A**) Heatmap representation of cold-evoked CT in mutant GAL4 control and GAL4-UAS-mediated rescue conditions, where FP is flaccid paralysis and CT is contraction. N=120, n=30 for each condition. (**B**) % of animals from rescue experiments performing a particular CT response, where NR is No Response (<10% reduction in length), Weak is ≥10% reduction in length, Moderate is ≥20% reduction in length, and Strong is ≥30% reduction in length; bars show proportions in % ± SEP.

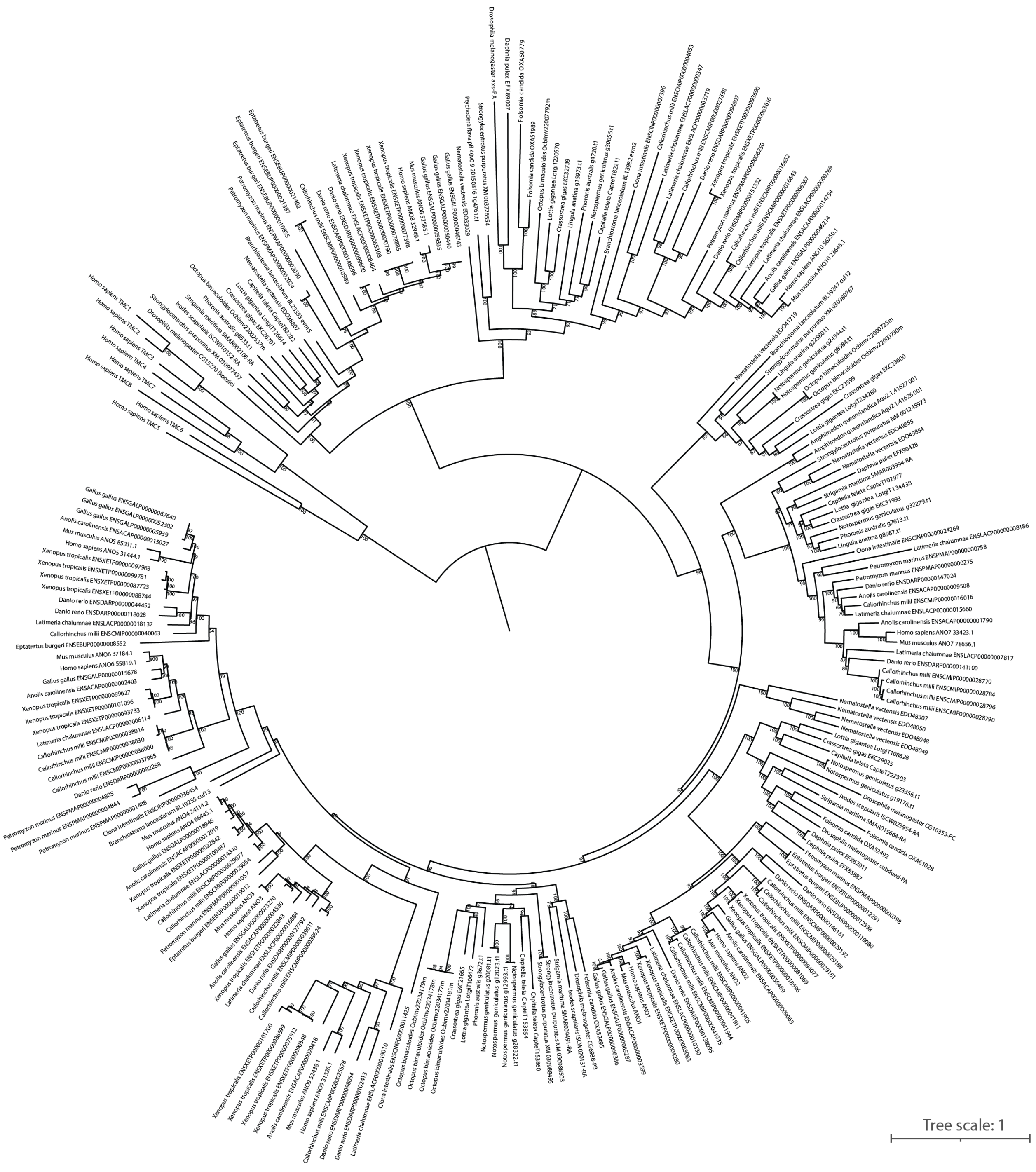


**Figure S6.** Maximum likelihood phylogeny of anoctamins corresponding to Figure 4.

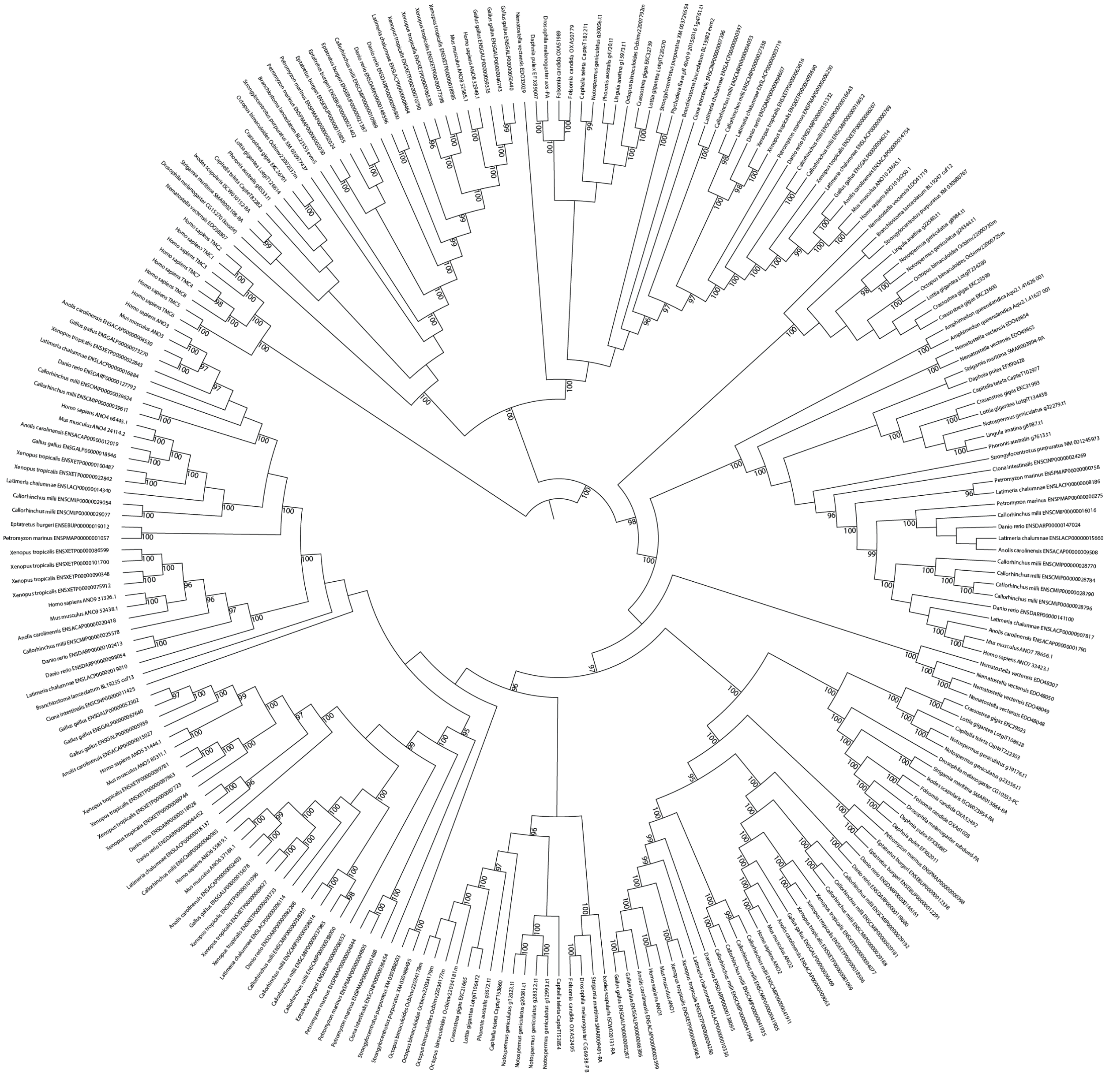

**Figure S7.** Rectified and rearranged phylogeny of anoctamins corresponding to Figure 4.

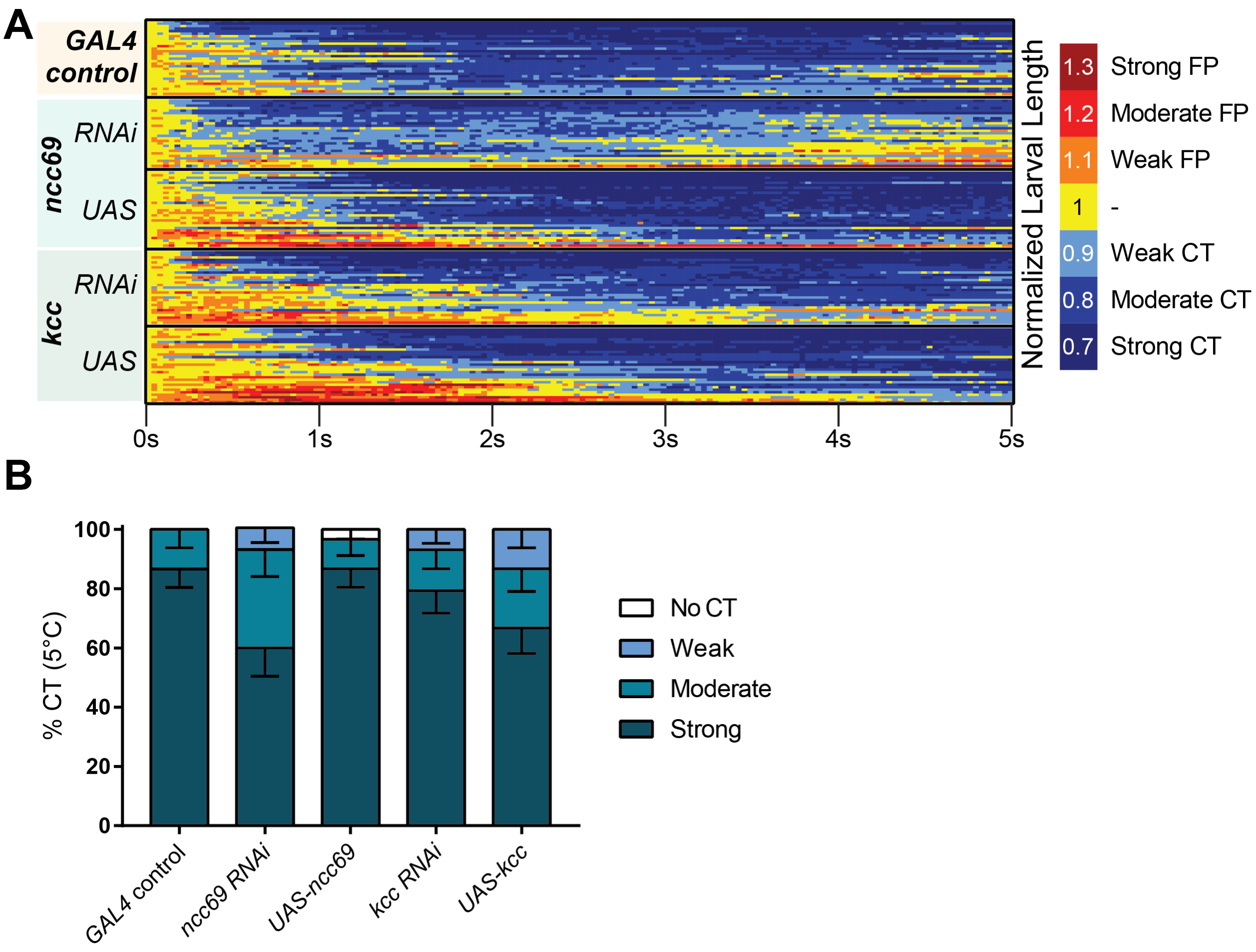


**Figure S8.** (**A**) Heatmap representation of cold-evoked CT in genetic control (*GAL4*), knockdown, and overexpression conditions, where FP is flaccid paralysis and CT is contraction. N=148; control n=30, *ncc69 RNAi* n=29, *ncc69 OE* n=30, *kcc RNAi* n=29, *kcc OE* n=30.=. (**B**) % of animals from SLC12 manipulation experiments performing a particular CT response, where NR is No Response (<10% reduction in length), Weak is ≥10% reduction in length, Moderate is ≥20% reduction in length, and Strong is ≥30% reduction in length; bars show proportions in % ± SEP.

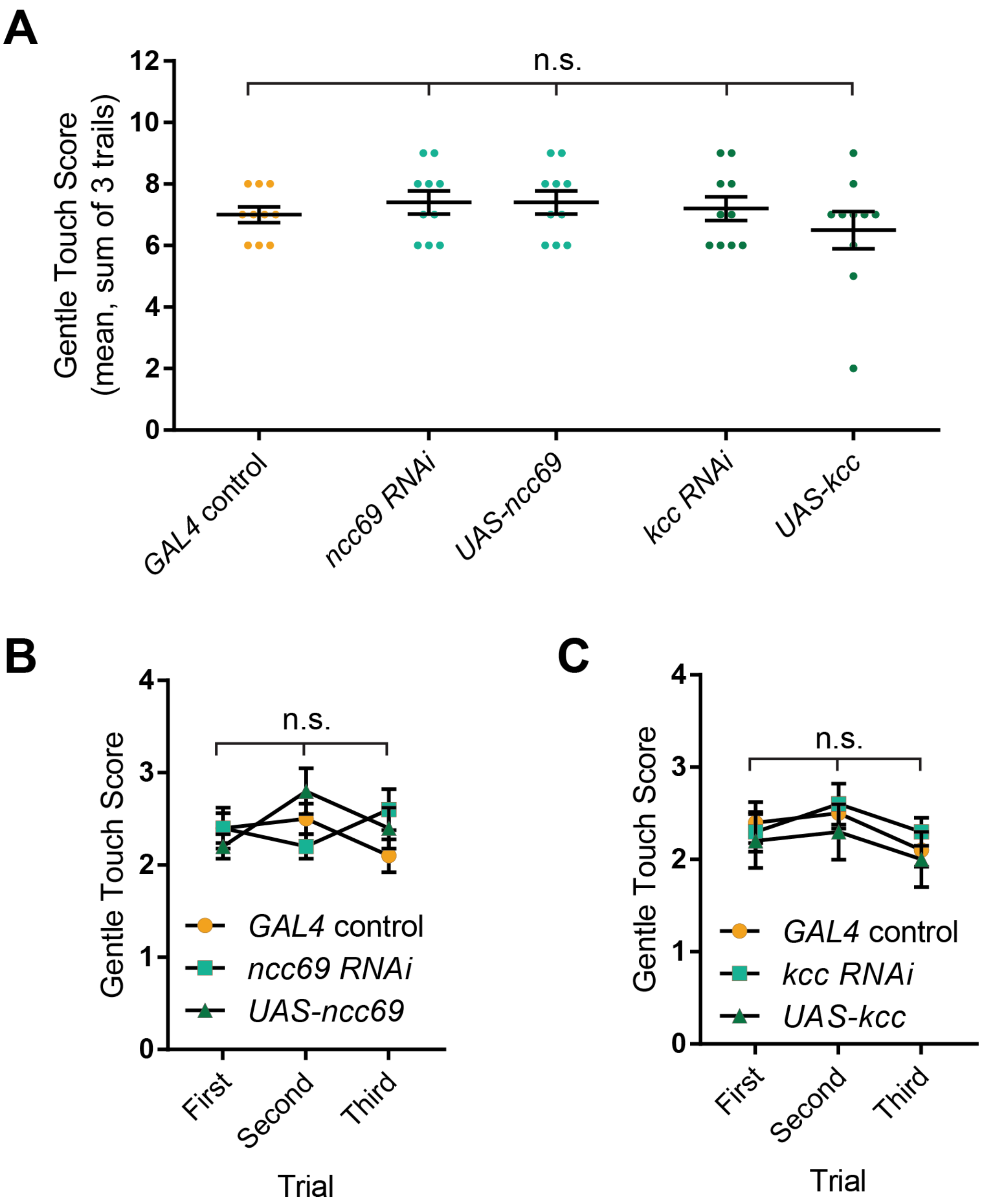


**Figure S9.** Results of gentle touch assay under SLC12 manipulations N=50, n=10 for each condition. Manipulation of SLC12 cotransporter expression does not affect gentle touch mechanosensitivity. (**A**) Summed touch scores, as in Figure 4A. (**B**) Average scores across trials for *ncc69* manipulations, as in Figure 4B. (**C**) Average scores across trials for *kcc* manipulations, as in Figure 4B.

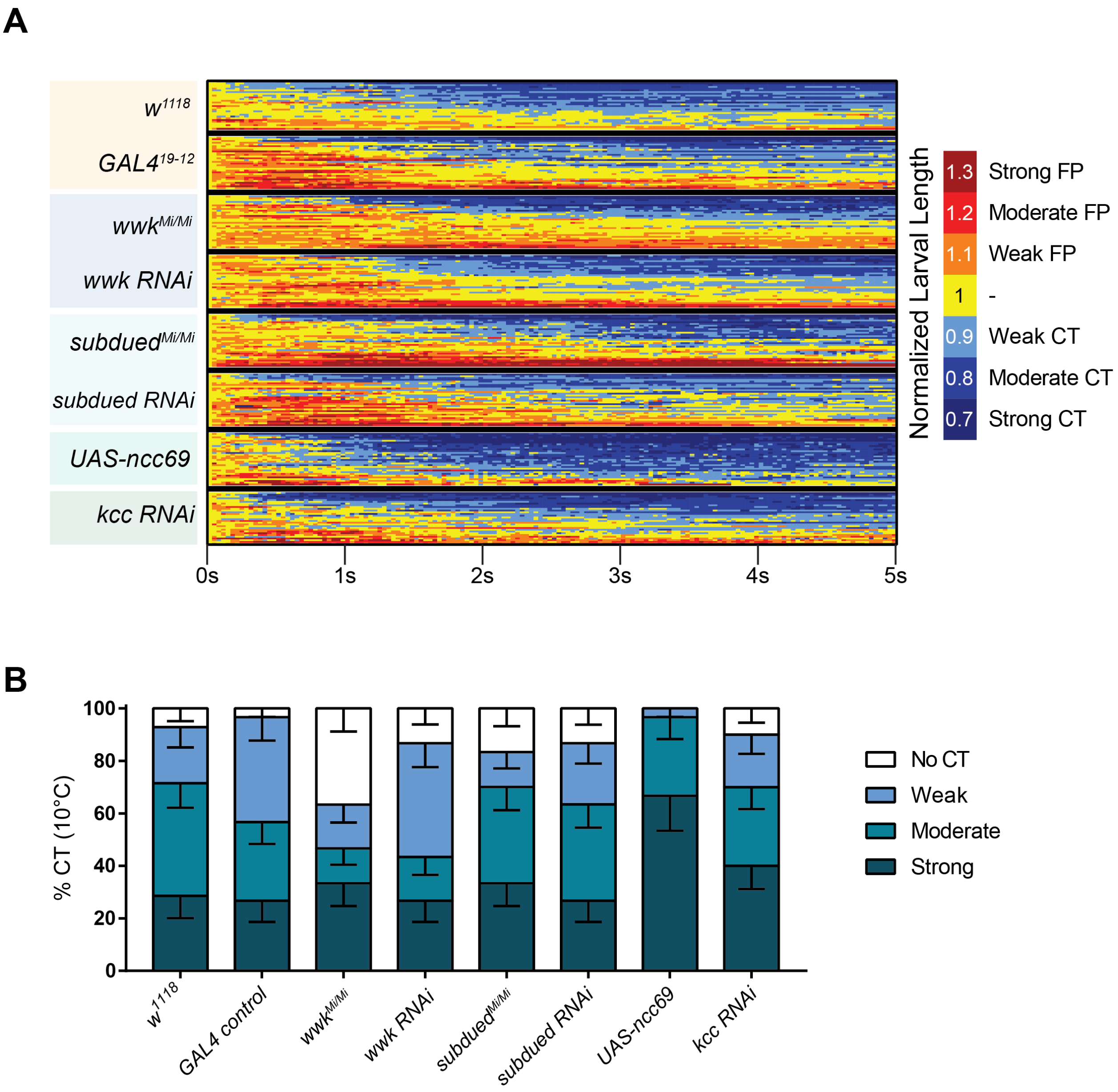


**Figure S10.** (**A**) Heatmap representation of cold-evoked CT in genetic control (*$w^{1118}$* or *$GAL4^{19\text{-}12}$*), knockdown, and overexpression conditions, where FP is flaccid paralysis and CT is contraction. N=239, n=30 for each condition except *$w^{1118}$*, where n=28. (**B**) % of animals from SLC12 manipulation experiments performing a particular CT response, where NR is No Response (<10% reduction in length), Weak is ≥10% reduction in length, Moderate is ≥20% reduction in length, and Strong is ≥30% reduction in length; bars show proportions in % ± SEP.
